# Genetic Impacts on Variability of Body Fat Distribution Uncover Gene-Environment and Gene-Gene Interactions

**DOI:** 10.64898/2026.03.31.715615

**Authors:** Xiaopu Zhang, Roby Joehanes, Jiantao Ma, Oliver Pain, Daniel Levy, Kenneth E. Westerman, Jordana T Bell

**Author notes:** Correspondence: Xiaopu Zhang, Jordana T Bell.

## Abstract

Body fat distribution is a strong predictor of cardiometabolic disease risk. Gene-environment and gene-gene interactions can affect body fat distribution, resulting in differential phenotypic variance across genotype groups that can be detected through variance quantitative trait loci (vQTLs). Using UK Biobank MRI data in conjunction with genetic data, we explored evidence for vQTLs for body fat distribution phenotypes aiming to uncover potential genetic interactions. We identified three vQTLs for liver fat distribution, including rs738408 (*PNPLA3*), rs4293458 (*APOE*), and rs58542926 (*TM6SF2*), and one vQTL region (*FTO*) for abdominal subcutaneous adipose tissue. To dissect putative gene-environment and gene-gene interactions underlying these signals, we identified multiple vQTL-environment interactions and one epistatic effect (rs58542926*rs429358) for liver fat. The vQTLs and interaction results were validated in multiple UK Biobank and external replication cohort datasets (Framingham Heart Study, All of Us, and TwinsUK), showing replication of the three liver vQTLs with the greatest reproducibility for vQTL rs738408. Our findings uncover vQTLs and underlying interaction effects on body fat distribution, especially liver fat, that may be useful for the development of precision medicine approaches.

## Introduction

Obesity is a major global public health challenge because it increases the risk of insulin resistance and metabolic syndrome, leading to cardiometabolic disease and its associated clinical complications. Body mass index (BMI) is a widely used indicator of obesity. However, several studies have shown that body fat distribution, especially intra-abdominal fat composition, is a more accurate measure of obesity and predictor of cardiometabolic disease risk than BMI [1], [2]. Although waist-to-hip ratio (WHR) can approximate intra-abdominal fat, imaging techniques such as magnetic resonance imaging (MRI) provide more precise assessments of body fat distribution [3].

Both genetic architecture and environmental factors play crucial roles in determining body fat distribution. For example, genome-wide association studies (GWAS) have established rs738409 (I148M) in *PNPLA3* as a strong genetic signal for liver fat [4], while genetic variants in the *FTO* gene region are associated with visceral [5] and subcutaneous adipose tissue accumulation [6]. Beyond genetic impacts, body fat distribution also varies with age and sex; and regular exercise and a healthy diet can reduce body fat volume and change body fat distribution [7], [8], [9], [10]. Genetic variants and environmental factors can also interact to influence body fat distribution, resulting in gene–environment (GxE) and gene–gene (GxG) interactions. GxG interactions have rarely been explored in the context of body fat distribution, but several studies have identified SNP-SNP interactions affecting BMI [11], [12]. On the other hand, there are several examples of GxE effects on cardiometabolic phenotypes. For example, *PNPLA3* 148M carriers have an elevated risk of metabolic dysfunction–associated steatotic liver disease (MASLD), which is driven by fat accumulation [13]. However, these individuals also respond to lifestyle changes to reduce intrahepatic triglyceride content, total cholesterol, and low-density lipoprotein cholesterol, in contrast with *PNPLA3* 148I homozygous individuals [14]. Such GxE interactions suggest a potential to modulate obesity-related disease risk in specific allele carriers through targeted lifestyle interventions. Therefore, capturing genetic interaction effects is essential for providing personalised cardiometabolic disease risk management.

Several studies have explored GxE and GxG interactions through genome-wide interaction studies (GWIS) [15], [16]. However, GWIS require specification of predefined environmental factors and are not ideal for comprehensive GxG analyses because of the heavy computational burden [15], [16]. In comparison with single GWAS, GWIS have a much greater multiple testing burden, affecting power. To address the limitations of GWIS, detecting genetic effects on phenotype variability, or variance quantitative trait loci (vQTLs), is an alternative approach to efficiently capture interaction effects [17], [18]. The principle of the vQTL is that if one allele is more sensitive to environmental changes or genomic context than the other alleles at a locus, this would result in different phenotypic variability across genotypes [17]. Therefore, vQTLs can capture genetic variants involved in GxE and GxG interaction effects. In this case rather than undertaking a full-scale GWIS, interaction analyses can only be performed on vQTLs rather than the genome-wide, which is a more efficient and powerful approach to detect interactions compared to GWIS.

Several studies have identified vQTLs for obesity-related phenotypes in large-scale cohorts [18], [19], [20], [21], [22]. For example, Wang et al (2019) [18] identified 75 vQTLs across 9 traits, including waist circumference, hip circumference, WHR, and BMI, in European-ancestry participants. Similarly, Lu et al (2022) [23] identified 185 vQTLs for 15 quantitative traits in the UK Biobank, 9 of which are related to obesity. Westerman et al (2022) [21] performed a vQTL meta-analysis across multiple ancestries, identifying 182 vQTL-phenotype associations and 132 independent vQTL-environment interactions related to plasma proteins and lipid traits. Variants in the *FTO* gene region have consistently been identified as vQTLs for BMI [18], [19], [20], [23], [24]. However, no study to date has identified vQTLs for imaging-measured body fat distribution.

To fill this knowledge gap, we conducted vQTL analyses of approximately 50,000 samples using UK Biobank genotype data in conjunction with magnetic resonance imaging data [25]. We explored genetic effects on phenotypic variance underlying eight body fat composition phenotypes in the UK Biobank dataset. The traits included four MRI-measured phenotypes including abdominal subcutaneous adipose tissue (ASAT), visceral adipose tissue (VAT), liver proton density fat fraction (PDFF), and pancreas PDFF, as well as four indices calculated from MRI measures including abdominal fat ratio, total abdominal adipose tissue index, total trunk fat volume, and weight-to-muscle ratio (Figure 1). We detected three vQTLs for liver PDFF (rs738408, rs429358, rs58542926) and one vQTL region (rs1285330517) for ASAT, which subsequent analyses show are involved in both GxE and GxG interactions. Further validation and replication analyses of additional UK Biobank data and in external cohorts (FHS, All of Us and TwinsUK) demonstrate the robust nature of these effects.

**Figure 1.**
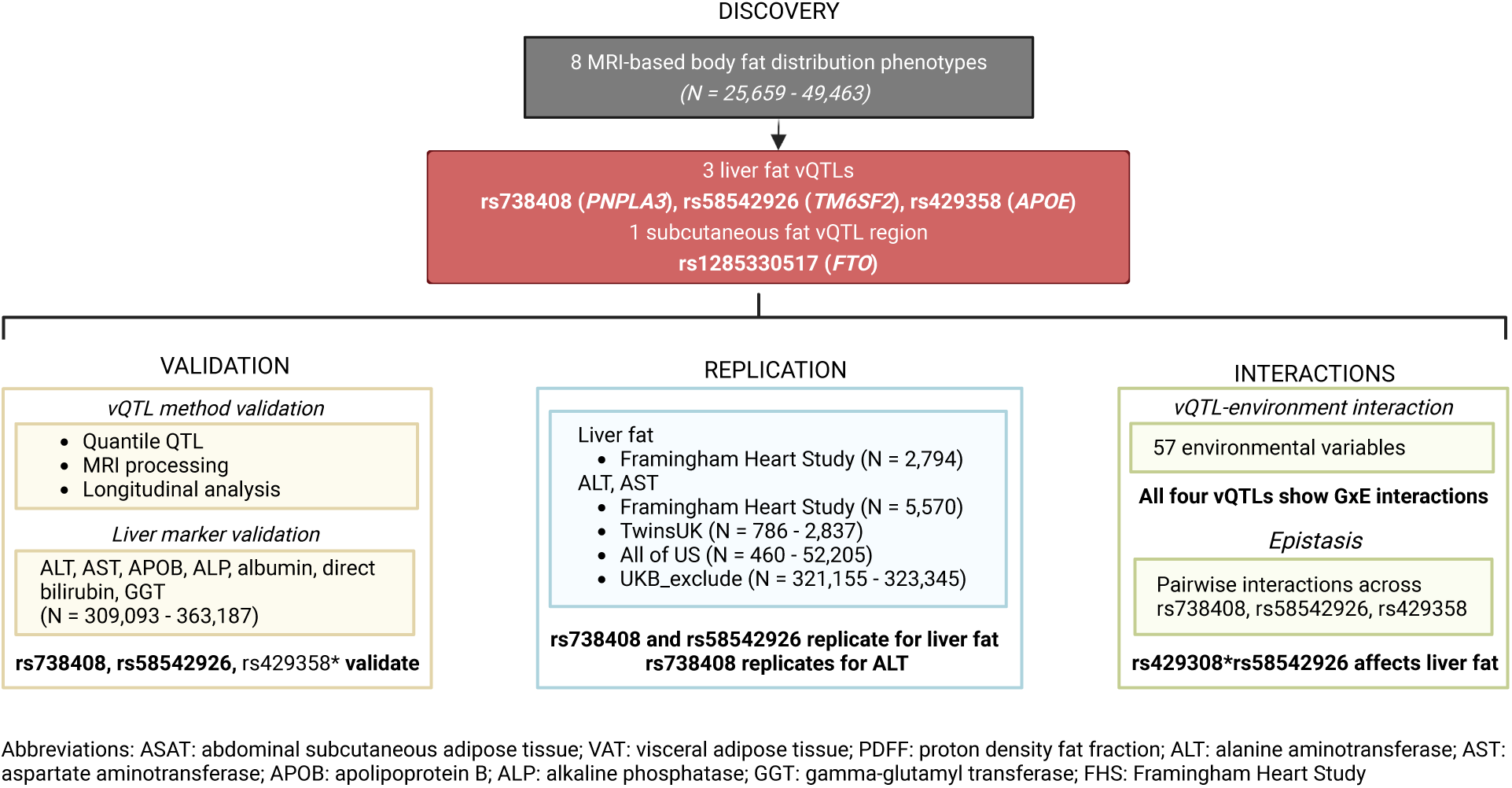
Study workflow. vQTLs were discovered using UK Biobank MRI datasets. After identifying four vQTL regions, validation analyses were performed within the UK Biobank using additional datasets. Replication was conducted in external cohorts. vQTLs interaction analyses were conducted including vQTL-environment and epistasis interactions. *rs429358 was not nominally significant for longitudinal stability in a data subset.

## Results

### Variance QTL analyses of body fat distribution imaging phenotypes in UK Biobank

We explored UK Biobank MRI phenotypes related to body fat distribution, focusing on phenotypes with more than 20,000 samples profiled to ensure robust detection of effects. As a result, four MRI-measured phenotypes including abdominal subcutaneous adipose tissue (ASAT), visceral adipose tissue (VAT), pancreas proton density fat fraction (PDFF), and liver PDFF, as well as four phenotype-derived indices including abdominal fat ratio, total abdominal adipose tissue index, total trunk fat volume and weight-to-muscle ratio, were retained for analysis (see Methods, Table 1, Figure 1). Given that individuals with the same BMI may vary in body fat distribution [26], we detected vQTLs for the eight traits both with and without adjusting BMI effects (see Methods).

**Table 1.** UK Biobank samples phenotype information. The information is summarized on the retained samples after data processing. The Field ID is derived from the UK Biobank dataset.

| Phenotypes (Field ID) | Sample size (samples with BMI) | % Female | Age range (median age) |
| --- | --- | --- | --- |
| ***Discovery set*** |  |  |  |
| ASAT (22408) | 49,449 (48,032) | 51.5 | 44-85 (65) |
| VAT (22407) | 49,463 (48,015) | 51.5 | 44-85 (65) |
| Liver PDFF (24352) | 41,534 (40,144) | 51.4 | 44-85 (66) |
| Pancreas PDFF (21090) | 26,445 (25,659) | 51.3 | 44-85 (65) |
| Abdominal fat ratio (22434) | 35,105 (33,995) | 52 | 45-82 (64) |
| Total trunk fat volume (22410) | 35,711 (34,592) | 51.6 | 45-82 (64) |
| Total abdominal adipose tissue index (22432) | 35,711 (34,592) | 51.6 | 45-82 (64) |
| Weight-to-muscle ratio (22433) | 35,138 (34,027) | 52.1 | 45-82 (64) |
| ***Validation set*** |  |  |  |
| ASAT (21086) | 34,788 (33,833) | 51.6 | 44-85 (65) |
| liver PDFF (21088) | 27,209 (26,294) | 51.3 | 46-82 (65) |
| liver PDFF (40061) | 36,170 (34,919) | 52 | 45-82 (65) |
| APOB (30640) | 361,394 (360,231) | 53.9 | 38-74 (58) |
| ALT (30620) | 363,037 (361,870) | 53.9 | 38-73 (58) |
| AST (30650) | 360,663 (361,825) | 53.9 | 38-73 (58) |
| ALP (30610) | 363,187 (362,017) | 53.8 | 38-73 (58) |
| Albumin (30600) | 332,625 (331,543) | 53.4 | 38-73 (58) |
| Direct bilirubin (30660) | 309,093 (308,162) | 49.4 | 38-73 (58) |
| GGT (30730) | 362,989 (361,821) | 53.8 | 38-73 (58) |

Genome-wide vQTL detection included three approaches (Brown-Forsythe (BF) test, deviation regression model (DRM), and squared residual value linear model (SVLM)), previously shown to have low false positive and relatively high discovery rates in simulation scenarios [17]. We also identified mean-controlling QTLs for these phenotypes using REGENIE [27]. Because the phenotypes are correlated, we estimated the effective number of traits using a previously described method [18], resulting in an effective number of 4.1 phenotypes, and a study-wise Bonferroni threshold of P<1.2e-8 (P<5e-8/4.1). Genetic variants surpassing Bonferroni threshold in at least one vQTL method and surpassing P<5e-8 in the other two vQTL mapping methods were considered vQTLs (Figure 1), whereas variants surpassing P<1.2e-8 in REGENIE were considered mean-controlling QTLs. Several quality control steps were applied to minimise false positive findings due to mean-variance associations and to retain conditionally independent vQTLs (see Methods).

Following genome-wide vQTL analysis of the eight MRI-derived body fat distribution phenotypes in 25,659-49,463 samples from the UK Biobank, significance threshold assessment across multiple vQTL detection methods, and quality control checks, there were altogether four vQTL regions detected (Figure 1, Table 2, Supplementary Figure 1A). Three vQTL regions identified by all three vQTL identification methods were associated with liver PDFF variability - rs738408 (*PNPLA3*) was a vQTL for liverPDFF after BMI adjustment (liverPDFF.adjBMI), rs58542926 (*TM6SF2*) was a vQTL for liver fat without adjusting for BMI, and rs429358 (*APOE*) was a liver PDFF vQTL regardless of BMI adjustment (Table 2). All three liver PDFF vQTLs were previously reported as mean-controlling GWAS signals for hepatic fat or liver function disorder [14], [28], [29]. The lead vQTLs of ASAT with BMI adjustment (ASAT.adjBMI) identified by SVLM (rs1285330517), DRM (rs62048402), and BF (rs11642015) methods are different, but these three lead vQTLs were located within the same linkage disequilibrium (LD) block in the *FTO* gene region, which was previously associated with obesity [30], [31]. We selected rs1285330517 as the lead vQTL for ASAT.adjBMI in subsequent analyses because it has the most significant P-value across the three lead vQTLs (P=3.19e-10).

**Table 2.** vQTL regions for liver PDFF and ASAT. RA refers to the reference allele and AA refers to the alternative allele (effect allele).

| **vQTLs** | **RA** | **AA** | **Gene** | **MRI trait** | **P(BF)** | **P(DRM)** | **P(SVLM)** |
| --- | --- | --- | --- | --- | --- | --- | --- |
| rs738408 | C | T | PNPLA3 | Liver PDFF.adjBMI | 2.28e-76 | 2.36e-75 | 2.25e-46 |
| rs429358 | T | C | APOE | Liver PDFF.adjBMI | 4.61e-24 | 4.47e-25 | 7.84e-17 |
| rs429358 | T | C | APOE | Liver PDFF | 1.28e-31 | 8.19e-33 | 7.45e-19 |
| rs58542926 | C | T | TM6SF2 | Liver PDFF | 2.74e-104 | 8.32e-103 | 4.34e-47 |
| rs1285330517 | CAAA | C | FTO | ASAT.adjBMI | 9.67e-09 | 2.21e-08 | 3.19e-10 |
| rs62048402 | G | A | FTO | ASAT.adjBMI | 2.15e-09 | 1.05e-08 | 3.24e-10 |
| rs11642015 | C | T | FTO | ASAT.adjBMI | 2.06e-09 | 1.06e-08 | 3.28e-10 |

All four vQTL regions identified in our study were previously identified either as vQTLs or as being in LD with vQTLs, but for different metabolic traits [18], [21], [23]. The vQTL region in the *FTO* gene was previously associated with variability in BMI, body fat percentage, waist circumference, hip circumference, and basal metabolic rate [18]. rs738408 (*PNPLA3*) was detected as a vQTL for alanine aminotransferase (ALT) and triglycerides (TG), and was in close LD with a vQTL for aspartate transaminase (AST) [21]. rs429358 (*APOE*) was a vQTL of ALT [21] and low-density lipoprotein (LDL) [23], and vQTL rs58542926 (*TM6SF2*) was associated with TG variability [23].

### vQTL effects validation within additional UK Biobank datasets

To minimize potential false positive findings, we implemented two vQTL validation steps within UK Biobank datasets (see Methods, Figure 1). First, we assessed whether vQTL variants manifested as quantile QTLs, which is an alternative measure of association with phenotypic variability. Next, we validated vQTL effects using several other datasets within UK Biobank. These included validation analysis of vQTL effects using MRI datasets processed by different institutes and longitudinal validation of vQTL effects in approximately 2,000 individuals with baseline visits and follow-up visits after 1 to 9 years.

Quantile QTLs estimate genetic effects in different quantile-based phenotype groups [32]. If the genetic effects remain consistent across quantile groups, the genetic variant likely affects only the mean level of the phenotype. Conversely, if genetic effects differ across phenotype quantiles, the variant manifests as a quantile QTL, and is therefore associated with phenotypic variability [32]. We categorized samples into ten groups according to their phenotype quantiles and estimated genetic effects in the first (Q1), fifth (Q5), and tenth (Q10) quantile groups. As a result, all four vQTL regions were identified as quantile QTLs (Supplementary Figure 1B, Supplementary Table 1). The effect of vQTL rs738408 (*PNPLA3*) was significant in Q5 and Q10 for liver PDFF (P<0.05), but not in Q1. vQTLs rs58542926 (*TM6SF2*) and rs429358 (*APOE*) significantly influenced Q10 group of liver PDFF (P<0.05), but not Q1 and Q5. The ASAT lead vQTL rs1285330517 (*FTO*) had significant effects in both Q1 and Q10 with opposite directions (P<0.05, C allele carriers have lower ASAT in Q1, but higher ASAT in Q10; Supplementary Figure 1B, Supplementary Table 1). As a result, we validated that the four vQTLs are related to phenotypic variability using an alternative approach.

Given that MRI scans were processed by three institutions in the UK Biobank using different pipelines (see Methods), we then validated vQTLs using further imaging datasets to confirm the robustness of vQTLs to processing methods (Figure 1). In addition to the discovery dataset, there were two additional liver PDFF datasets (Field ID 21088 and 40061) and one additional ASAT dataset (Field ID 21086) with smaller sample sizes (Table 1, see Methods). Using samples included in all datasets (28,948 in three liver PDFF datasets and 38,882 in both ASAT datasets), we observed high correlations across the three liver PDFF datasets (Spearman’s correlations range from 0.98 to 0.99) and between the two ASAT datasets (Spearman’s correlation is 0.99). All four vQTL regions validated using these three datasets (P<0.05, Supplementary Table 2) and the three liver PDFF vQTLs also validated at the Bonferroni threshold used in the discovery stage as well (P<1.2e-8, Supplementary Table 2).

To explore the longitudinal stability of the detected vQTL effects, we analysed individuals with two MRI visits in the UK Biobank (Figure 1). For the tested phenotypes, altogether 2,076 to 2,380 participants had a follow-up MRI scan 1 to 9 years after baseline scan (see Methods). We conducted vQTL analysis at baseline and at follow-up in this smaller subset of participants. The three liver PDFF vQTLs (rs738408, rs4293458, and rs58542926) surpassed the nominal significance threshold in the first timepoint dataset, and rs738408 and rs58542926 also validated in the follow-up visit dataset at nominal significance (Supplementary Table 3). However, rs429358 did not validate in follow-up visit data, and the ASAT vQTL was not significant at both timepoints (Supplementary Table 3). which could be because the lower sample size did not provide sufficient power to detect effects. To investigate this, we randomly selected 2,000 samples from the discovery set and evaluated whether the BF, DRM, and SVLM methods could detect vQTLs at a nominal significance threshold (P<0.05) in this smaller sample size (see Methods, Supplementary Table 4). We repeated this subsampling procedure 100 times to calculate the statistical power for vQTL detection. As a result, all three methods reached 80% power for vQTL rs58542926. For rs738408, BF and DRM reached 80% power, whereas SVLM achieved 77% power. The power to detect vQTLs rs1285330517 ranged from 12% to 15%, and power to detect vQTL rs429358 ranged from 51% to 74%, across the three methods. When the subset size was increased to 2,500, power to detect rs738408 and rs58542926 increased to 94%. The power to detect rs1285330517 and rs429358 increased slightly, but remained below 80%, except for DRM when testing rs429358 (Supplementary Table 4). Therefore, the results from these analyses suggest that the effects of liver fat vQTLs rs738408 and rs58542926 are longitudinally stable, albeit in a sample subset. However, the smaller sample size for these longitudinal analyses has limited the power to validate vQTLs rs429358 and rs1285330517.

Overall, we observed that the identified vQTLs for liver fat exhibit consistent effects when applying a different approach to detect genetic effects on phenotypic variance, and they are highly robust across imaging data processing methods, and most exhibit longitudinal stability.

### vQTL-Environment interactions underlying vQTLs

To assess whether the detected vQTLs could capture genetic interaction effects, we next carried out vQTL-environment factor interaction analyses. Following a comprehensive search of the UK Biobank dataset, 57 environmental factor variables were selected for interaction analysis (see Methods, Supplementary Table 5). The 57 variables capture information related to diet (23 of 57 variables), alcohol use, demographic information and lifestyle factors such as smoking, sleep, sedentary behavior, and physical activity. We undertook vQTL-environment interaction analyses of these data, using three approaches to define the “environmental factor” (Figure 2A).

**Figure 2.**
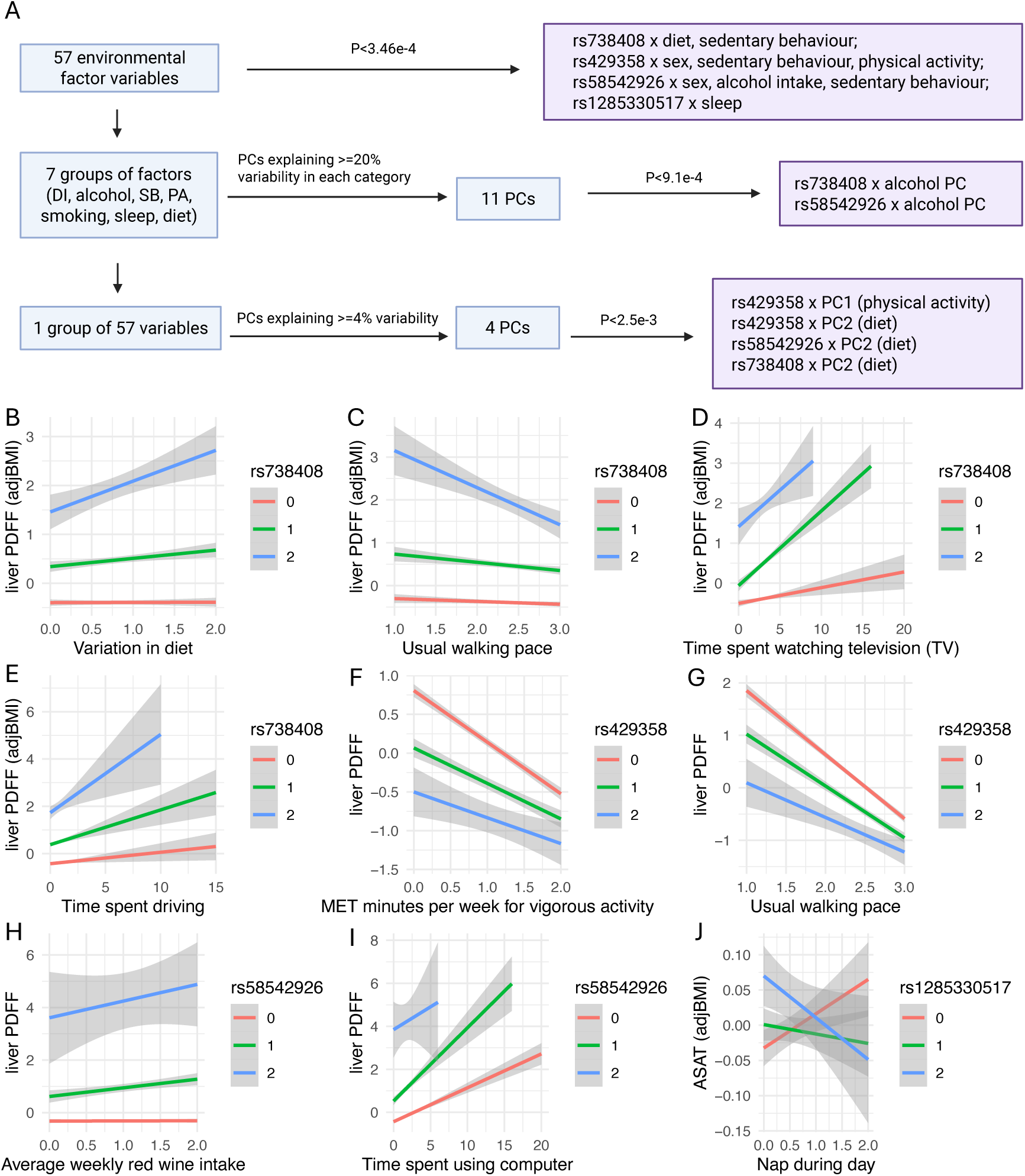
vQTL-environment interaction analysis. A) Summary of three strategies for vQTL-environment interaction analyses. B-J) Results of interactions between vQTLs and individual factor variables. Plots show the direction and size of genetic effects on the phenotype with a change in environmental factor. Genotype 0 represents the homozygous group for the reference allele, while genotype 2 represents the homozygous category for the alternative allele. The reference/alternative alleles are as follows: rs738408 (C/T), rs429358 (T/C), rs58542926 (C/T), and rs1285330517 (CAAA/C). Y-axis shows the residuals of each phenotype after adjusting the covariates.

We first considered vQTL-environment factor interaction analyses where the environmental factor was each of the 57 variables. Because these variables are correlated, we estimated the effective number of factors to be 28.9 (see Methods), and applied a Bonferroni multiple testing correction for the interaction analysis (P<0.05/5x28.9). All three liver PDFF vQTLs, rs738408, rs429358, and rs58542926, exhibit interaction effects with variables that capture aspects of physical activity (PA) and sedentary behaviour (SB). The effect of variant rs738408 (*PNPLA3*) on liver fat changed with SB and PA variables related to time spent watching television, driving, as well as usual walking pace, but also frequency of dietary change (Figure 2B-E). The effect of variant rs429358 (*APOE*) on liver fat showed dependence on PA variables related to usual walking pace and vigorous activity (Figure 2F-G). Lastly, the effect of rs58542926 (*TM6SF2*) on liver PDFF was affected by SB variable ‘time spent using the computer’, as well as average weekly red wine intake (Figure 2H-I). Specifically, these three liver fat vQTL variants were previously reported as GWAS signals for hepatic fat or liver function [14], [28], [29], and we observed that each environmental factor had a consistently more pronounced effect on the GWAS risk allele (T of rs738408, T of rs429358, T of rs58542926) for liver fat or liver function, compared to the non-risk allele. Lastly, the ASAT lead vQTL, rs1285330517 (*FTO*), showed an interaction with the frequency of taking nap during the day (Figure 2J).

Furthermore, age and sex were also included in these 57 variables. Given that body fat distribution varies across sex and age groups [7], [8], [33]. we also explored interaction effects between vQTLs and sex or age. As a result, liver fat vQTLs rs429358 (*APOE*) and rs58542926 (*TM6SF2*) exhibited significant interactions with sex (Supplementary Figure 2, Supplementary Table 6), indicating these two SNPs show sex-specific effects on the population-level liver fat volume, which are consistent with the sex differences in the risk of fatty liver diseases [34]. The risk allele of rs58542926 was found to have greater effects on liver PDFF in men than in women [35]. rs429358 risk allele carriers have higher TC and LDL levels and the effect is more significant in women than men with Alzheimer’s disease [36]. We further explored the potential for sex-specific interaction effects by investigating second-order interaction factors (G-sex-E). This analysis yielded 62 significant interaction effects (P<0.05/(5x28.9), Supplementary Table 7), suggesting that additional interactions could be uncovered through sex-stratified analysis. However, as the primary aim of the current study is to characterize interactions across the entire cohort regardless of sex, we did not pursue these second-order interactions further.

Because some of these 57 environmental factor variables are related to each other, we used two additional approaches to categorise environmental factors in exploring interactions. The second vQTL-environment factor interaction analysis considered groups of related environmental factor variables. Here, the 57 environmental factor variables were grouped into 7 categories including demographic information (DI), diet, alcohol consumption, smoking, sleep, sedentary behavior (SB), and physical activity (PA) (Supplementary Table 5). We then carried out a principal component analysis (PCA) of the environmental factor variables within each of the 7 categories, and retained principal components within each group that explained more than 20% of the variance (Figure 2A). As a result, the first PC for DI, diet, PA, smoking, the first two PCs for SB and sleep, and the first three PCs for alcohol consumption were retained. We then carried out vQTL-environment factor analyses where the environmental factor was each retained PC, for altogether 11 PCs tested. At a Bonferroni threshold (P<0.05/(5x11)), interaction effects were observed between liver PDFF vQTLs rs738408 (*PNPLA3*) and rs58542926 (*TM6SF2*) with the second PC for the number of glasses of alcohol intake weekly (Supplementary Table 6), which was primarily correlated with spirits intake (Pearson correlation r2=0.70). When using a nominal threshold (P<0.05), seven more interactions were observed between three liver fat vQTLs and PCs for alcohol consumption, PA, sleeping, and smoking.

The final vQTL-environment factor interaction analysis considered all 57 environmental factor variables as an entire group (Figure 2A). Similarly, we performed PCA across all 57 environmental factor variables and selected the first four PCs that captured more than 4% variability across all 57 variables. We then carried out vQTL-environment factor analyses with each retained PC as the environmental factor, for altogether 4 PCs tested. Four vQTL-PC interactions for liver PDFF surpassed Bonferroni correction (P<0.05/(5x4)). Specifically, vQTL rs429358 (*APOE*) interacted with PC1, which was strongly correlated with variables related to PA. vQTLs rs429358 (*APOE*), rs58542926 (*TM6SF2*), and rs738408 (*PNPLA3*) showed significant interaction effects with PC2, which was primarily associated with diet-related factors (Supplementary Table 6). No more interactions were observed when using a nominal significance threshold.

Overall, we used three different approaches to categorise environmental factors in undertaking vQTL-environmental factor interactions that may underlie the observed vQTL signals. All three liver PDFF vQTLs (rs429358, rs58542926, and rs738408) showed interactions with environmental factors in at least two of the approaches. Two liver PDFF vQTLs (rs58542926 (*TM6SF2*) and rs738408 (*PNPLA3*)) showed interactions with environmental factors in each of the three approaches, but the specific interacting factors varied depending on how the environmental factors were defined. A limitation in this analysis was the difference in sample sizes across the three approaches, which leads to different power, because samples with missing data were excluded in the second and third approaches, which apply PCA. Moreover, 23 of the 57 variables related to diet, leading to diet-related PCs being over-represented in the final approach when all variables were included in the PCA.

### Validation of liver fat vQTLs and GxE interactions using liver function markers

Because liver fat accumulation leads to liver dysfunction, we further explored whether the liver PDFF vQTLs and their GxE interaction effects also affect serum liver function markers in a larger UK Biobank dataset (N = 309,093-363,187; Table 1; part of the sample overlaps with the discovery UK Biobank MRI sample). Liver function phenotypes focused on alanine aminotransferase (ALT) and aspartate aminotransferase (AST). We also considered additional phenotypes related to liver function for this validation analysis, including apolipoprotein B (APOB), alkaline phosphatase (ALP), albumin, direct bilirubin, and gamma-glutamyl transferase (GGT). All three liver PDFF vQTLs (rs738408, rs58542926, and rs4293458) affected the variability of ALT, AST, ALP, albumin, and direct bilirubin at the Bonferroni threshold (P<0.05/3, Supplementary Table 8). Furthermore, rs58542926 and rs4293458 were also vQTLs for GGT and APOB (P<0.017=0.05/3), but rs738408 was not. Besides, rs58542926 manifested as a vQTL for albumin (P<0.017), but rs429358 and rs738408 did not.

We also validated the liver fat GxE interaction effects using the seven liver function phenotypes. Using the same threshold as the previous interaction analysis (P<3.5e-4), 7 of 10 GxE interactions for liver PDFF were validated in at least one of the seven liver function markers (Supplementary Table 9). Specifically, 5 of 7 interactions with sex, time spent watching television, time spent driving, variation in diet, and usual walking pace, were associated with ALT levels, which is the most clinically relevant marker for liver function disorders.

In conclusion, our analyses indicate that liver fat vQTLs and their potential underlying GxE interaction effects could be validated using liver function markers.

### Replication of liver fat vQTLs

To assess the reproducibility of the liver fat vQTLs, replication analyses were conducted in three external cohort datasets, including TwinsUK, All of Us, and the Framingham Heart Study (FHS), as well as in UK Biobank datasets that excluded individuals with MRI scans in the discovery set (UKB_exclude) (see Methods, Figure 1). We first sought replication of the three liver PDFF vQTLs (rs58542926 in *TM6SF2*, rs4293458 in *APOE*, and rs738408 in *PNPLA3*) in the FHS dataset using CT-measured liver fat deposition. We also aimed to replicate these three liver PDFF vQTLs using liver function biomarkers ALT and AST in all replication datasets (TwinsUK, All of Us, FHS, and UKB_exclude).

We first carried out replication analysis of the three liver PDFF vQTLs using liver fat deposition phenotypes in the FHS dataset (N = 2,794), which were assessed using CT scans rather than MRI scans as in the discovery dataset. As a result, vQTLs rs738408 (P<2.3e-7) and rs58542926 (P<4.8e-3) showed significant effects at a Bonferroni correction threshold (P<0.017=0.05/3), whereas vQTL rs429358 did not replicate (Table 3, Supplementary Table 10). This could be due to limited statistical power attributed to the small replication sample size. Our power calculations with a sample size of 2,500 show that the BF and SVLM methods have lower than 80% power to detect rs429358, although DRM reaches 80% power (see Methods, Supplementary Table 4). The power of BF increases to 80% when the sample size increases to 3,000. Another limitation of the small sample size is that the dataset may be too small to adequately capture a wide range of phenotype variability.

**Table 3.**
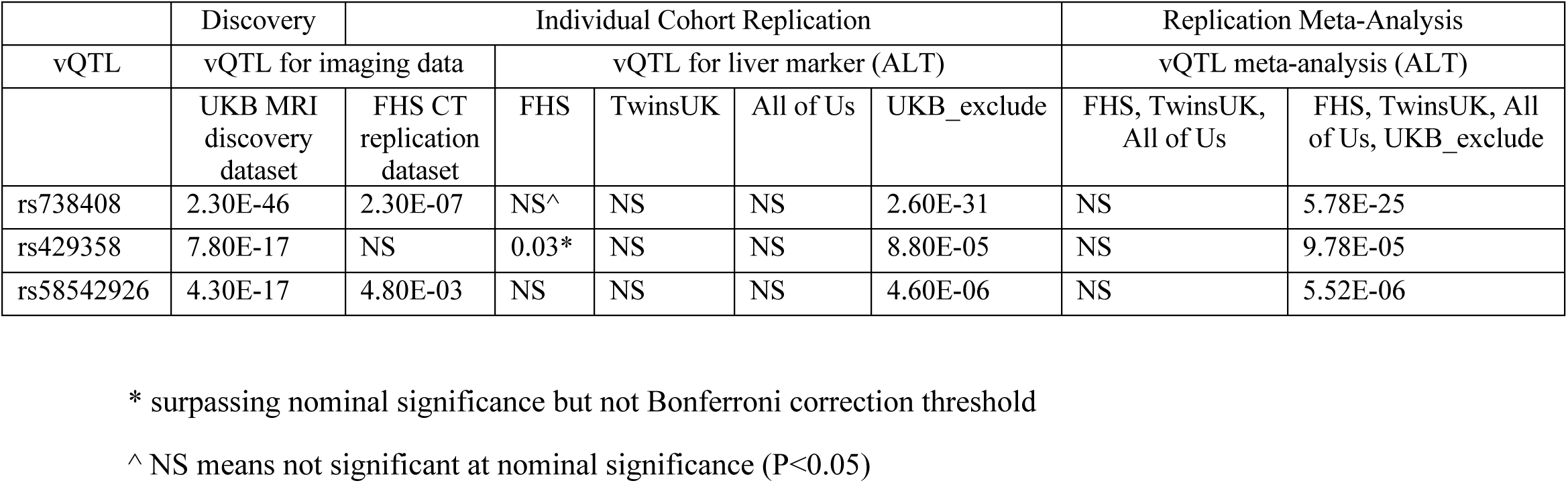
Strength of association for liver PDFF vQTLs in discovery, validation, and replication datasets. For each vQTL, BF, DRM and SVLM were applied to test its significance with and without adjusting for BMI. The p-value shown in the table is the least significant p-value, eg. 2.30E-46 is the largest p-value observed when conducting vQTL analysis using BF, DRM, and SVLM methods to test rs728308 using UKB liver PDFF dataset (including liver PDFF.adjBMI and liver PDFF without adjusting BMI), the p-values observed in the remaining tests are more significant than 2.30e-46 (p<2.30e-46).

We next sought replication of the three liver PDFF vQTLs using liver function markers ALT and AST measured in All of Us (N = 460–52,205), FHS (N = 5,570), TwinsUK (N = 786–2,837), and UKB_exclude (N = 321,155–323,345). All three vQTLs replicated as vQTLs for ALT and AST in UKB_exclude at the Bonferroni correction threshold (P<0.017), and rs738408 (*PNPLA3)* showed vQTL effects for ALT in all four datasets at nominal significance using at least one vQTL detection methods (Table 3, Supplementary Table 11). We then conducted meta-analysis of the vQTL results across these four replication cohorts (see Methods). Using summary statistics from the DRM and BF, a meta-analysis across the four replication cohorts identified all three genetic variants, rs429358, rs738408, and rs58542926, as vQTLs for ALT and AST at the Bonferroni correction threshold (P<0.017). Meta-analysis using SVLM summary statistics similarly confirmed all three as vQTLs for ALT, although only rs738408 was associated with AST (P<0.017). Given that UKB_exclude has a substantially larger sample size than the other datasets, it may strongly influence the meta-analysis results, therefore, we also conducted a meta-analysis restricted to All of Us, FHS, and TwinsUK. Using DRM and BF summary statistics, rs738408 was confirmed as a vQTL for ALT and AST (P<0.017). With SVLM summary statistics, none of the vQTLs as detected from the meta-analysis at the Bonferroni threshold, but the rs738408 was detected as a vQTL for ALT at the nominal significance threshold (P=0.048) (Supplementary Table 12).

Therefore, the replication analyses provide support for the reproducibility of the liver PDFF vQTLs. vQTLs rs738408 (*PNPLA3*), rs58542926 (*TM6SF2*), and rs429358 (*APOE*) showed consistent associations with liver biomarkers (ALT and AST) in several external datasets, with rs738408 demonstrating the most reproducible effects across all cohorts.

### vQTLs uncover epistasis for liver fat

vQTLs uncover not only interactions with environmental factors, but also epistatic effects. Given that we observed three vQTLs associated with liver fat, we explored pairwise interaction effects between the three vQTLs, rs58542926 (*TM6SF2*), rs4293458 (*APOE*), and rs738408 (*PNPLA3*), on liver PDFF. As a result, we detected an interaction interaction surpassing Bonferroni correction threshold (P<0.017=0.05/3) between vQTL rs58542926 in *TM6SF2* and vQTL rs429358 in *APOE* (P=4e-5) (Supplementary Figure 3), but no significant interactions were detected for the other pairs of tested vQTLs. We examined the epistasis effect in two additional liver fat datasets processed by different methods within UK Biobank (Field ID 21088 and 40061, see Methods) previously used in vQTL validation. This sensitivity analysis showed that the epistatic effect was validated in both datasets (P=3.6e-4 in Field ID 21088 dataset, P=6.1e-4 in Field ID 40061 dataset).

We further explored whether the identified epistatic effect (rs58542926 and rs429358) could also influence liver function biomarkers within UK Biobank. We conducted the interaction analysis between rs58542926 and rs429358 using the 7 liver marker factors in UK Biobank (N = 309,093-363,187, Table 1, part of the sample overlaps with the discovery UK Biobank MRI sample, see Methods). At the Bonferroni threshold (P<7e-3=0.05/7), the epistatic effect was validated for ALT (P=2.5e-5 for ALT, P=1.2e-5 for ALT.adjBMI), AST (P=1.2e-3 for AST, P=1.2e-3 for AST.adjBMI), ALP (P=9.7e-5 for ALP, P=1.1e-4 for ALP.adjBMI), APOB (P=4e- 15 for APOB, P=2.7e-15 for APOB.adjBMI), and direct bilirubin levels (P=5.2e-4 for bilirubin, P=6e-4 for bilirubin.adjBMI).

We then sought replication of the epistatic interaction in the four replication datasets, starting with CT-measured liver fat deposition phenotype in FHS (N=2,794). The epistatic interaction did not replicate in the FHS liver fat dataset at a nominal significance threshold (Supplementary Table 10), although the direction of the interaction effect is as expected. To assess whether this was attributable to limited statistical power, we performed 100 iterations of random sampling, selecting 3,000 samples from the discovery set to estimate the frequency with which interaction effects surpassed the nominal significance threshold (see Methods). Our results indicate that the statistical power to detect gene-gene interactions in this sample size is 20% (Supplementary Table 4). Therefore, the failure to validate the epistasis effect is likely due to the insufficient sample size. We then examined whether this interaction can be detected for ALT or AST in the four replication datasets. Except for UKB_exclude, no replication of the epistatic effect was observed in the individual cohort, FHS, All of Us, and TwinsUK (Supplementary Table 11). The epistatic interaction reached significance (P<0.025=0.05/2) for both ALT and AST in a meta-analysis across the four replication datasets (Supplementary Table 12). However, a meta-analysis of the epistatic effects in the three replication datasets excluding UKB_exclude was not significant. In summary, evidence for the epistatic interaction was weak outside of the UKB discovery datasets and the UKB_exclude dataset, potentially due to the smaller sample size and limited power to detect the interaction effect.

## Discussion

The current study identified four unique vQTLs for MRI-derived measures of liver PDFF and ASAT in the UK Biobank. Several validation analyses within UK Biobank datasets show that these vQTLs exhibit very robust effects. Replication of two liver PDFF vQTLs, rs738408 and rs58542926, using CT-measured liver fat deposition, and replication of all three liver PDFF vQTLs using liver function biomarkers ALT and AST in external cohorts, confirm the robust nature of these vQTL effects. Follow-up vQTL-environment and targeted vQTL-vQTL interaction analyses uncovered putative interplay between vQTLs and specific environmental factors, as well as gene-gene interactions between two liver fat vQTLs.

Previous studies have detected several vQTLs related to cardiometabolic risk factors, including vQTLs for anthropometric indices [18] and blood markers [21] of cardiometabolic health. These non-additive genetic effects on cardiometabolic profiles provide valuable insights into disease risk monitoring and potential avenues for targeted interventions. Compared to traditional metrics, for example BMI, body fat deposition measured by imaging techniques may serve as a more reliable predictor of disease. However, no prior vQTL studies have been conducted on imaging-derived body fat distribution. The current study addresses this gap.

Three vQTLs for liver PDFF and one vQTL region for ASAT were detected using the UK Biobank MRI scans. These vQTLs regions and nearest genes have all been previously linked to metabolic phenotypes in the literature. Specifically, all three liver PDFF vQTLs, rs58542926 (*TM6SF2*), rs738408 (*PNPLA3*), and rs429358 (*APOE*), were previously associated with liver lipidation [13], [28], [29], [30]. *TM6SF2* plays an important role in processing nascent very low-density lipoprotein (VLDL) particles into fully lipidated VLDLs, which transport lipids from the liver to the bloodstream to supply energy [28]. vQTL rs58542926 in *TM6SF2* is a nonsynonymous variant (E167K) that reduces TM6SF2 mRNA levels, impairing nascent VLDL particles processing and leading to lipid accumulation in the liver. vQTL rs429358 is a missense variant that defines the isoform APOE4 allele. Individuals carrying the APOE4 allele (rs429358-C) tend to have higher levels of VLDL particles [37], promoting lipid transportation from the liver to the bloodstream. vQTL rs738408 is in close LD with rs738409, which leads to the I148M variant of gene *PNPLA3*, a strong genetic risk factor of metabolic dysfunction–associated steatotic liver disease (MASLD) [13]. A recent study showed that the PNPLA3-I148M variant changes the morphology of the Golgi apparatus, which interacts with lipid droplets, leading to liver disease [38]. Lastly, the vQTL region associated with ASAT was in the *FTO* gene, where many previous studies have identified genetic risk variants for obesity across ancestries [30]. Several studies have found that *FTO* functions as a demethylase of N6-methyladenosine (m6A) and regulates adipogenesis in an m6A-dependent manner [31]. For example, *FTO* affects the m6A level around the splice site of adipogenic regulator *RUNX1T1* to regulate its alternative splicing [31]. The detection of these vQTLs was differentially impacted by BMI adjustment. rs429358 remained a significant vQTL regardless of BMI adjustment. rs738408 manifests as vQTL for liver fat and rs1285330517 is a vQTL for ASAT only after adjusting for BMI, suggesting that BMI-related changes in total body weight may otherwise mask these effects. In contrast, rs58542926 is a vQTL for liver fat without BMI adjustment, which implies a potential association with both variability of liver fat and BMI. These observations suggest that future vQTL studies of body fat distribution should evaluate vQTLs both with and without BMI adjustment to avoid missing context-specific genetic effects.

Waist-to-hip ratio (WHR) is an anthropometric measure related to body fat distribution. Compared to MRI-based measures of central adiposity, WHR is easier to obtain but is also less accurate in quantifying central body fat distribution. Previously, two large-scale vQTL studies observed several vQTLs for WHR. Wang et al (2019) [18] found rs459193 as a vQTL for WHR, but no vQTLs for WHR after BMI adjustment using around 348,000 samples. Lu et al (2022) [23] used a similar sample, but did not detect any vQTL for WHR. Despite the smaller sample size in the current study, we identified more vQTLs for body fat distribution. This is likely attributable to the more precise quantification of adiposity in MRI-derived phenotypes.

To explore how robust the detected vQTL effects are, we conducted replication analyses using CT-measured liver fat data in the FHS cohort. vQTLs rs738408 and rs58542926 replicated as vQTLs of liver fat deposition in FHS at a Bonferroni correction threshold, but vQTL rs429358 was not significant, which is likely at least in part attributed to limited power. We also explored replication using liver function biomarkers AST and ALT in the All of Us, TwinsUK, FHS, and UK Biobank subset which excludes MRI samples. rs738408 (*PNPLA3)* was the most replicated vQTL signal for ALT and AST, both from each cohort-specific dataset and meta-analysis results. As a result, rs738408 (*PNPLA3)* is the most robust vQTL obtained in the current study.

We observed that the vQTL meta-analysis results based on the BF and DRM statistics identified more vQTLs than those based on the SVLM statistics. For example, rs738408 was detected as a vQTL for ALT at Bonferroni threshold in the meta-analysis of FHS, TwinsUK, and All of Us using DRM and BF statistics, but not when using SVLM statistics. However, the effect of rs738408 on ALT surpassed nominal significance (P<0.05) in the SVLM meta-analysis. The reason could be due to differences in input data and methodological frameworks of the three approaches. DRM and BF calculate the distance of each observation from the median value in each genotype group and subsequently apply either a linear regression model (DRM) or an ANOVA test (BF) to assess the association between genotypes and the distances. SVLM first regresses the phenotype on genotype to obtain residuals and then uses the squared residuals as the dependent variable in a linear regression model with genotype as the predictor. Therefore, DRM and BF may provide more similar meta-analysis results, while the lower discovery rate of SVLM may arise from its methodological design. This observation aligns with a previous simulation study illustrating that BF and DRM perform similarly across various scenarios [17]. Besides, BF and DRM show higher discovery rates than SVLM when the phenotype follows a non-Normal distribution [17], which is relevant here as liver fat distribution is skewed.

Three of our identified liver fat vQTLs were already established GWAS signals of liver fat, with previously established main effects on the phenotype [5]. On the other hand, there were several other liver fat GWAS main effect signals that we did not identify as vQTLs. To investigate this, we used the summary statistics from a previous study by Liu et al (2021) [5], which performed a GWAS on MRI-measured liver fat in the UK Biobank using a similar sample size. Liu et al (2021) [5] identified 12 GWAS signals for liver fat, 3 of which are our identified liver fat vQTLs (rs429358, rs58542926, rs738408). Six of the remaining main effect liver fat GWAS signals (rs1723945191, rs188247550, rs2642438, rs4665985, rs55714539, rs7029757) reached nominal significance (P<0.05) as vQTLs across the BF, DRM, and SVLM results, but failed to meet the stringent vQTL filtering and multiple-testing threshold. The remaining 3 main effect liver fat GWAS variants (rs112875651, rs113070129, rs1233772390) did not surpass nominal significance in any vQTL identification method. This can be attributed to several factors. First, the power to identify vQTL is lower than power to detect mean-controlling QTLs at the same sample size. Second, the majority of genetic effects on phenotypes are likely to be additive, with fewer vQTLs expected compared to additive effect QTLs [39], [40].

To explore whether the vQTL results could capture genetic interactions, we carried out a systematic vQTL-environment analysis using three approaches. Altogether, 11 vQTL-environment interactions were observed when testing each environment factor variable individually. Liver PDFF vQTLs were involved in the majority of interactions, showing strong interactions with sex, sedentary behavior, and physical activity measures. Subsequently, we also grouped environmental factor variables into several categories or considered them all collectively as one group. Because the individual variable factors are correlated with each other, these categorizations aim to assess whether vQTLs are more likely to interact with variables related to a particular category. We conducted PCA in both groups and selected the top PCs to represent each of the 7 categories, as well as the whole variable group. The results differed from those obtained when analysing each factor variable individually. For example, the vQTLs-diet interactions were not found when examining individual dietary factors, except for the interaction between rs738408 and variation in diet. However, diet had significant effects when analyzing PCs across all variables related to the diet category. Several factors may explain the discrepancies across the three vQTL-environment interaction analyses. First, the statistical power of the three approaches differs due to the variation in sample size. Samples with missing exposure data were excluded in the PCA. Therefore, PCs of the whole exposure variable list (57) were based on a much smaller sample size (N = 16,735–19,972), compared to the analyses involving PCs of individual categories (N = 28,742–47,720), and those of vQTL by each exposure variable interaction (N = 28,899–47,984). Second, some categories may be over-represented in the PCA because of differences in the number of exposure variables within each category. For example, 23 of the 57 factor variables were diet-related, which may explain why the analysis highlighted interactions with diet when considering all 57 variables as one group.

Previous research on GxE interactions aligns with our identified vQTL-environment interaction results. We observed that three liver fat vQTLs (rs738408 in *PNPLA3*, rs429358 in *APOE*, and rs58542926 in *TM6SF2*) are more likely to interact with physical activity (PA) and sedentary behavior (SB) when considering the environmental factors individually. Previous studies have found that increasing physical activity could mitigate the risks associated with the rs429358 by decreasing the VLDL particle sizes [29]. The effect of rs738409, which is in close LD with rs738408, on fatty liver and liver disease also depends on exercise intensity [41]. In contrast, few studies have reported the interaction effect between variants in *TM6SF2* and physical activity levels on liver PDFF. Our finding of the interaction between rs58542926 and PA warrants further exploration of the mechanisms underlying liver fat accumulation and may help inform future clinical trials. Additionally, the interaction effect that we observed between rs1285330517 (*FTO*) and the frequency of taking a nap on ASAT is similar to findings from Young et al [42], who identified that rs1421085 in gene *FTO* interacts with deviation from mean sleep duration and influences obesity. When categorizing the 57 variable factors into different groups, we obtained the interactions between two liver fat vQTLs (rs738408 and rs58542926) and alcohol intake, as well as interactions between all three liver vQTLs and diet. Previous findings support the observation that heavy alcohol use increases the risk of liver diseases in individuals who are rs58542926 and rs738408 risk allele carriers [43]. Similarly, dietary intake has been shown to modulate the genetic effects of *APOE, TM6SF2*, and *PNPLA3* on liver fat accumulation. For example, omega-3 fatty acid intake has been found to reduce the detrimental effects of *APOE* rs429358 of hepatic fat by lowering the synthesis of new VLDL particles and triglycerides from the liver [29]. Another study identified that rs58542926 (*TM6SF2*) interacts with multiple dietary factors, including vegetable, fruit, and legume intake, with T-allele carriers exhibiting an amplified dietary effect on hepatic function [44]. The detected vQTL rs738408 (*PNPLA3*) is in close LD with rs738409 (PNPLA3-I148M variant). A previous study found that with a 500-kcal diet restriction, a much stronger reduction in hepatic lipids was seen in 148MM homozygotes than in 148II homozygotes [45].

We also identified a vQTL-vQTL epistatic interaction effect between rs429358 (*APOE*) and rs58542926 (*TM6SF2*) on liver PDFF, which has not been previously reported to our knowledge. The T alleles of rs429358 and rs58542926 are the two GWAS risk alleles for hepatic fat [3]. We observed that the effect of rs58542926 T allele on liver PDFF depends on rs429358 genotype. Specifically, individuals carrying the rs58542926 T allele exhibit higher liver PDFF when they are also rs429358 T allele carriers compared to rs429358 C allele carriers. This interaction is biologically plausible, and existing evidence offers indirect support for it. Specifically, vQTL rs429358 in *APOE* has been associated with APOB levels [46], and *TM6SF2* has been shown to interact with and stabilise APOB in mouse models [47]. Furthermore, variant rs58542926 in *TM6SF2* has been demonstrated to reduce APOB secretion in human hepatic 3D spheroids [48]. Together, these findings suggest functional links between genes *TM6SF2* and *APOE*, and the observed epistatic effect provides additional motivation for further investigation of the interplay between these two genetic loci. Although the epistatic effect did not replicate using CT-measures of liver fat, it was shown to influence liver function markers in the UKB_exclude data subset, as well as across the four replication cohorts, but not in meta-analyses excluding UKB samples. One reason why our analyses did not show replication of the epistatic effect outside of UK Biobank might be that the sample size is limited, therefore affecting power to detect epistasis.

The current study has several other limitations. First, although UK Biobank has collected extensive questionnaire data that are valuable for exploring vQTL-environment interactions, the majority of these data were not collected at the same time as the MRI scan. Therefore, we only considered environmental exposure variables collected at the same time as the MRI scan data, which limited the number of environmental factor variables and sample size. Second, the serum liver markers were collected at a different timepoint from the MRI images obtained by the UK Biobank. Although we validated the vQTL effects, vQTL-environment interactions, and the epistatic effect on liver function markers, the comparability would be improved if these liver function biomarker and liver imaging datasets were collected contemporaneously and from the same participants. Third, the sample sizes for the replication analysis are limited to conduct a powerful replication study, and especially for epistasis. The power of the liver fat epistasis test conducted in FHS (N = 2,794 participants) is much smaller compared to the test carried out in the UK Biobank discovery dataset (N = 41,534). Similarly, we replicated the episttic effect on ALT and AST in approximately 340,755 samples in UK Biobank, but failed to replicate this in external cohort datasets of a few thousand samples. Fourth, the current study only focused on vQTLs of MRI-measured body fat distribution in European individuals because the number of individuals from other ancestries with MRI data is not sufficient to conduct a powerful vQTL analysis.

Using vQTLs as indices to uncover interaction effects is an efficient approach to identify if the genetic effect on body fat distribution depends on context, and this could benefit the development of precision medicine approaches. For example, the current study shows that rs738408 and rs58542926 show interaction with weekly alcohol intake to influence liver fat. Risk-allele carriers for these variants may derive greater benefit from reducing alcohol consumption to decrease liver fat, compared to rs429348 carriers. There are several future directions that can be explored based on this work. First, if sufficient samples from non-European populations become available, a multiple-ancestry replication of vQTLs and their associated interaction effects could be prioritised. Second, validation of the interaction effects *in vitro* and in clinical trials will be essential to explore the translational relevance of our findings, and may provide novel insights for health intervention.

In conclusion, the current study identified three vQTLs for liver fat and one vQTL region for ASAT. Follow-up vQTL–environment interaction and epistasis analyses confirmed that these signals may capture biologically meaningful GxE and GxG interactions. Validation within the UK Biobank and replication in external cohorts further demonstrated the robust nature of the three liver vQTLs, with rs738408 (*PNPLA3*) showing most reproducible effects across all datasets.

## Methods

### UK Biobank data

The UK Biobank is a long-term prospective cohort study [49]. UK Biobank collected biological samples and health-related records from over 500,000 individuals aged from 40 to 69 throughout the UK. Since 2014, approximately 85,000 volunteers have contributed to the imaging project, which collects scans from multiple tissues.

### Phenotypes selection

UK Biobank used magnetic resonance imaging (MRI) technique to capture the whole-body imaging, including the abdominal, brain and heart. We searched through the abdominal MRI datasets (Category ID 105) to filter the body fat distribution-related phenotypes and only kept those with more than 20,000 participants. As a result, eight phenotypes remained (Table 1). Four of them were MRI-derived body fat volumes: 1/ abdominal subcutaneous adipose tissue (ASAT), which measures the subcutaneous adipose tissue in the abdomen from the top of the femoral head to the top of the thoracic vertebrae T9, 2/ visceral adipose tissue (VAT), which measures the adipose tissue within the abdominal cavity, excluding adipose tissue outside the abdominal skeletal muscles, and adipose tissue and lipids within the cavity and posterior of the spine and back muscles, 3/ liver proton density fat fraction (PDFF), which is a fat referenced measurement that measures the average PDFF in up to nine (and at least three) regions of interest (ROI) in the liver; where the ROI avoid regions show inhomogeneities to ensure the accuracy, as well as major vessels, and bile ducts, and 4/ pancreas PDFF, which measures the pancreas fat fraction. The other four were indices from MRI-derived measures: 1/ abdominal fat ratio, or the total abdominal fat volume divided by the total abdominal fat volume and thigh muscle volume; calculated as 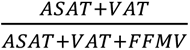, where FFMV represents the total thigh fat free muscle volume, 2/ total trunk fat volume, which is calculated as the total volume of VAT and ASAT, 3/ total abdominal adipose tissue index, which is the total abdominal fat divided by squared height, or 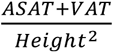, and 4/ weight-to-muscle ratio, an index of body weight divided by total thigh muscle volume.

Because abdominal adiposity phenotypes were derived by multiple institutes (Table 1), ASAT and VAT were processed by two different institutes (Field ID 22408 and 21086 for ASAT, Field ID 22407 and 21085 for VAT), and liver PDFF were supplied by three centres (Field ID 24352, 21088, and 40061). The samples for the same phenotype in different datasets largely overlapped, but the processing pipelines used by each institute may lead to differences. Therefore, for each phenotype, we selected the datasets with the largest sample size as the discovery set and the remaining datasets as validation sets. For example, liver PDFF with field ID 24352 (N = 49,449) was used as the discovery set while liver PDFF with field ID 21088 (N = 27,209) and field ID 40061 (N = 36,170) were used as validation sets. Besides, a subset of volunteers (N = 2,076-2,380) has participated in a follow-up imaging visit. This subset of longitudinal samples was used as validation as well.

Because liver fat accumulation leads to liver dysfunction, we also validated liver PDFF vQTLs in liver function markers. Liver function serum markers were collected from blood samples in a larger sample size (N=309,093-363,037) and measured by Beckman Coulter AU5800 chemistry analyzer. These markers included APOB (Field ID 30640), ALT (Field ID 30620), AST (Field ID 30650), ALP (Field ID 30610), albumin (Field ID 30600), direct bilirubin (Field ID 30660), and GGT (Field ID 30730) (Table 1).

### Samples level quality control filters

Due to the majority of volunteers in imaging projects being self-reported Europeans, we focused on genetically identified European-ancestry individuals in the current study. To define the ancestry groups, we used principal components analysis (PCA) to compare the UK Biobank individuals to the samples with known ancestries in 1000 Genome (1KG) project following the GenoPred pipeline [50]. Specifically, we performed PCA on linkage disequilibrium (LD)-pruned genetic variants from the 1KG samples and used the first six principal components (PCs) to train a multiclass classification model to distinguish individuals across five ancestry groups: Africans (AFR), Admixed Americans (AMR), East Asians (EAS), Europeans (EUR), and South Asians (SAS). The same genetic variants were used to calculate genetic PCs in the UK Biobank participants. The first six PCs from the UK Biobank were fit into the trained multiclass classification model to observe the predicted probability of each ancestry for all individuals. The individual is assigned to an ancestry group if the probability is higher than 50%. We compared the estimated ancestry with the estimated ethnic background in the UK Biobank (Field ID 22006). As a result, all estimated Caucasians were also estimated Europeans in our dataset, confirming the confidence of our estimated genetic ancestry of individuals. Besides, if there are any outliers within each population, they could bias the analysis. Therefore, we removed outliers within European individuals by performing PCA and using the distance from population centroids across 10 PCs. Outliers were defined as samples falling outside the threshold of the third quartile (75th percentile, Q3) plus 30 times the interquartile range 𝑄3 + 30 × 𝐼𝑄𝑅. We also removed samples with inconsistent self-reported sex and genetically estimated sex (Field ID 22001) and those suggested to be removed by the UK Biobank.

### Genotype data processing

Genotyping, imputation, and initial quality control of UK Biobank genotype data have previously been described in detail [49]. For samples in each phenotype dataset, we excluded genotype data with 1/ minor allele frequency less than 0.05, 2/ Hardy–Weinberg equilibrium (HWE) P-value less than 1e-6, 3/ individual missingness higher than 0.05, and 4/ imputation information less than 0.8. The final sample sizes for each phenotype with genotype data are listed in Table 1.

### Phenotype data processing

For each phenotype, we excluded the related samples with ukb_gen_samples_to_remove function in the R package ukbtools [51]. We also removed samples with phenotype values higher or lower than 5 standard deviations of the phenotype mean level. A linear mixed model was applied to adjust for covariates, including sex (Field ID 31), age (Field ID 21003), and the first 20 genetic PCs (Field ID 22009) as fixed effects, as well as assessment centre (Field ID 54), genotype array (Field ID 22000), and genotype array position (Field ID 22000) as random effects. Body mass index (BMI, Field ID 21001) was optionally included as a fixed effect along with the factors. Therefore, each phenotype had two input datasets for analysis: one without BMI adjustment (Equation 1) and one with BMI adjustment (Equation 2).

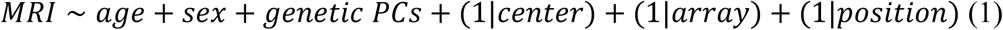

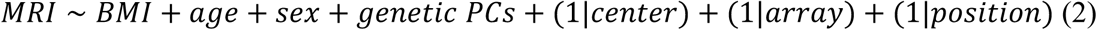

### Identification of GWAS signals and vQTLs

Based on results from a previous simulation study [17], we chose to apply three vQTL identification approaches, including the Brown-Forsythe (BF) test, deviation regression model (DRM), and squared variance linear model (SVLM), as the methods with low false positive rates. Given that these methods have different limitations, we applied all three methods in the analyses and only kept vQTLs that were detected by all methods [52]. We also carried out genome-wide association studies of these MRI traits using REGENIE to detect main effect QTLs [27]. Because of the correlation across these MRI measurements, we calculated the effective number of phenotypes as previously explained [53]. Briefly, if 𝜆_1_, …, 𝜆_*k*_ are the eigenvalues of the PCA, the ratio of the sum of eigenvalues to the sum of squared eigenvalues (Equation 3) indicates the independent dimensions captured by the PCs. Each eigenvalue reflects the variance explained by its corresponding PC. If all phenotypes are independent and contribute equally, each PC would have equal eigenvalues. However, if some PCs explain much more variance than others (i.e., eigenvalues are unequal), this ratio will be lower, indicating redundancy and fewer effective dimensions. Here, the independent dimensions correspond to the number of effective phenotypes (Equation 3).

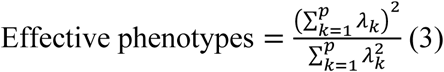

A genetic variant manifests as a vQTL if it 1/ surpasses the Bonferroni threshold in at least one vQTL detection method, and 2/ surpasses the threshold of P*<*5e-8 in the remaining two vQTL identification methods. A genetic variant that surpasses the Bonferroni threshold of the REGENIE result is classified as a mean-controlling signal.

To reduce the chance of false positive findings, we conducted several quality control steps. First, we carried out conditional analysis on vQTLs and mean-controlling QTLs for each phenotype with COJO [54]. Second, we performed LD clumping of conditionally independent vQTLs and mean-controlling signals for each phenotype using Plink [55], based on the P-values (LD window=500kb, 𝑟^4^=0.8). This step aimed to reduce false positive vQTLs arising from genetic variants that are in partial LD with mean-controlling signals, and vice versa [17]. Pruned signals were removed from either the vQTL or mean-controlling QTL list. Third, when a genetic variant was identified as both a vQTL and a mean-controlling QTL for the same phenotype, this genetic variant was taken as a covariate to adjust for its additive genetic effect on the phenotype (Equation 4). The residual values from Equation 4 and the genetic variant were refit into the vQTL identification method to evaluate whether the genetic variant still surpassed the significance threshold. This QC step eliminates false positives that may arise due to correlation between the mean level and variance of the data distribution [17].

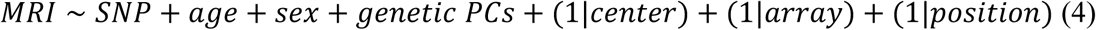

### Quantile QTLs

Quantile QTL was introduced as an approach to assess the heterogeneity of phenotypes across genotype groups [32]. To evaluate whether a genetic variant manifests as a quantile QTL, samples are divided into groups based on phenotype quantiles. The additive genetic effects of the genetic variant are estimated within each phenotype quantile group. Consistent genetic effects across all phenotype quantile groups suggest the absence of phenotype heterogeneity for genetic effects. Conversely, substantial variation in effect size across the phenotype quantiles indicates the presence of genetically associated phenotypic heterogeneity. To conduct quantile QTL analysis, we divided the phenotype into 10 quantile groups and estimated the genetic effects in the first (Q1), fifth (Q5), and tenth (Q10) quantiles with linear models using R in this study.

### Power analysis

To estimate vQTL detection power, we randomly selected 2,000, 2,500, and 3,000 samples from the liver PDFF and ASAT discovery datasets each. We then repeated the selection process 100 times and applied the BF, DRM, and SVLM methods to these data subsets. The discovery rate was defined as the frequency with which the results exceeded the nominal significance threshold (P<0.05). Under the assumption that the genetic variants manifest as vQTLs, this discovery rate is equivalent to the statistical power. Similarly, we performed 100 iterations of random selection (n=2,000, 2,500, and 3,000) within the discovery liver PDFF dataset to determine if the epistasis effect could be identified (P<0.05). This approach allowed us to estimate the empirical power to detect epistasis effects in datasets with different sample sizes.

### vQTL-environment interaction analysis

To comprehensively explore candidate environmental factors, we considered the entire UK Biobank dataset, which divided recorded information into 357 categories and each category contains the information fields. We excluded categories and fields that 1/ were not recorded during the same timepoint as the MRI imaging visit; 2/ could not represent long-term effects, for example, estimated nutrients consumed in a single day (Category ID 100098 in the UK Biobank dataset); and 3/ were unrelated to MRI measures, for example, biological sample processing information (Category ID 163). As a result, 83 environmental exposure fields remained, spanning information across lifestyle factors such as diet, physical activity, and sedentary behavior. Next, we used PHESANT [56] to process these 83 fields with default settings.

PHEASANT was developed for automated phenome scans in the UK Biobank to determine how to systematically process traits and conduct phenome-wide association studies. For different types of data, PHESANT applies different preprocessing steps. For example, if the information is discrete and more than 20% participants have the same value, PHESANT will remove values with fewer than 10 samples. After processing the 83 environmental exposure variables through PHESANT, we further excluded the environmental variables with more than 30% missing values across the 50,087 unique participants used for the MRI vQTL analysis. These exclusions resulted in a final set of 57 environmental exposure variables for follow-up interaction analysis.

For each phenotype, we adjusted the covariates as we did in the vQTL discovery step and then used the residuals as input for interaction analysis. For each vQTL-phenotype association, we first conducted interaction analysis with each environmental factor separately (vQTLxE) using Equation 5. Because these environmental factors are correlated, we also calculated the effective number of environmental factors using Equation 3. Subsequently, we also considered grouping the environmental factor data according to the type of environmental factors. The 57 environmental factors were manually categorized into seven groups as follows: demographic information (DI), diet, alcohol consumption, smoking, sleep, sedentary behavior (SB), and physical activity (PA) (Supplementary Table 5). PCA was applied to each category, and the PCs that explained at least 20% of the variance across the variables in each category were selected to represent that category. Lastly, we performed PCA on all 57 factors and selected the PCs which explained more than 4% of the total variance among these factors for interaction analysis.

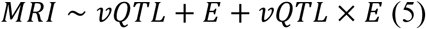

### vQTL-vQTL interaction analysis

vQTLs could capture epistasis effects (vQTL-vQTL interactions). Because there were three vQTLs for liver PDFF, we conducted pairwise interaction analysis across the three vQTLs (Equation 6). Phenotype processing followed the same procedure as the phenotype processing steps used for vQTL identification. We applied a linear model to fit the additive genetic effects of vQTL1 and vQTL2, as well as the interaction effects between vQTL1 and vQTL2 (vQTL1 x vQTL2). A statistically significant interaction effect (P<0.05) indicates that the interaction effect of vQTL1 and vQTL2 on MRI traits is independent of their additive effects, providing evidence of epistasis effect on the MRI traits.

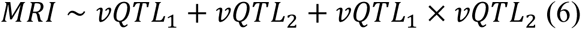

### Replication and Meta-analysis

We sought replication of vQTL effects and epistatic effects in three external cohorts, the Framingham Heart Study (FHS), All of Us, and TwinsUK (Figure 1).

FHS (https://dsgweb.wustl.edu/PROJECTS/MP1.html) is a multigenerational study which aims to identify risk factors for cardiovascular disease [57]. In this dataset, 2,794 participants had CT-measured liver fat deposition, while 5,563 to 5,570 samples had available ALT and AST levels from fasting blood samples for replication analyses [58]. Therefore, we conducted the replication of vQTLs and epistatic effects on both CT-measured liver fat deposition and liver markers in FHS.

The All of Us research program is a nationwide project which collected health information from individuals residing in the United States [59]. In this cohort, we carried out vQTL analysis and epistatic effect tests in 44,360 ALT and 49,961 AST “Caucasian” samples.

TwinsUK is the largest adult twin registry in the United Kingdom [60]. In TwinsUK, 1,243 and 2,837 participants had ALT and AST data from blood samples, respectively. The sample size decreased to 786 and 2,376 when restricting to samples with BMI data collected at the same timepoint. We used TwinsUK as the replication dataset for vQTLs and epistatic effects on ALT and AST.

We also conducted replication analysis of the vQTLs using ALT and AST in the extended UK Biobank dataset, which excluded participants with MRI scans used in the discovery dataset (UKB_exclude). As a result, there were 321,155 to 323,345 UKB samples with ALT and AST measures left in the replication dataset “UKB_exclude”.

The phenotype processing procedures for FHS, All of Us, TwinsUK and UKB_exclude datasets followed the same steps as those used in the UK Biobank discovery dataset. All three vQTL identification methods (BF, DRM, and SVLM) and the epistasis interaction model were performed in the replication cohorts.

We also conducted meta-analyses of the vQTL effects and epistasis for ALT and AST levels across the FHS, All of Us, TwinsUK, and UKB_exclude datasets, using a fixed-effect model in METAL based on P-values [61].

## Acknowledgements

We thank the research participants involved in the cohort studies included in this research manuscript. We thank Ting Qi and Jian Yang for implementing the vQTL methods in OSCA. We thank Tianyuan Lu for the insightful discussion and feedback for this project. XZ acknowledges the financial support of the China Scholarship Council. This project also received support from the JPI ERA-HDHL DIMENSION project and UK Biological Sciences Research Council (BBSRC, BB/S020845/1 and BB/T019980/1 to JTB), and in part by project BACMETH (selected funding by ERC, funded by EPSRC EP/Y023765/1 to JTB). OP is supported by the Wellcome Trust [222811/Z/21/Z].

TwinsUK is funded by the Wellcome Trust, Medical Research Council, Versus Arthritis, European Union Horizon 2020, Chronic Disease Research Foundation (CDRF), Wellcome Leap Dynamic Resilience Programme (co-funded by Temasek Trust), Zoe Ltd, the National Institute for Health and Care Research (NIHR) Clinical Research Network (CRN) and Biomedical Research Centre based at Guy’s and St Thomas’ NHS Foundation Trust in partnership with King’s College London.

The Framingham Heart Study was supported by NIH contracts N01-HC-25195, HHSN268201500001I, and 75N92019D00031.

Disclaimer: The views and opinions expressed in this manuscript are those of the authors and do not necessarily represent the views of the National Heart, Lung, and Blood Institute, the National Institutes of Health, or the U.S. Department of Health and Human Services.

## Author contributions

XZ and JTB designed the project. XZ led the data analysis. OP, KW, JM and RJ contributed to data analysis. XZ and JTB wrote the article.

## Materials & Correspondence

This research has been conducted using the UK Biobank Resource under application number 112124. Access to TwinsUK data can be applied through the cohort data access committee https://twinsuk.ac.uk/resources-for-researchers/access-our-data/.

## Reference

[1] S. Fujioka, Y. Matsuzawa, K. Tokunaga, and S. Tarui, “Contribution of intra-abdominal fat accumulation to the impairment of glucose and lipid metabolism in human obesity,” Metabolism, vol. 36, no. 1, pp. 54–59, 1987.

[2] I. Khan et al., “Surrogate adiposity markers and mortality,” JAMA Network Open, vol. 6, no. 9, pp. e2334836–e2334836, 2023.

[3] J. Linge et al., “Body composition profiling in the UK biobank imaging study,” Obesity, vol. 26, no. 11, pp. 1785–1795, 2018.

[4] S. L. Park et al., “Genome-wide association study of liver fat: The multiethnic cohort adiposity phenotype study,” Hepatology Communications, vol. 4, no. 8, pp. 1112–1123, 2020.

[5] Y. Liu et al., “Genetic architecture of 11 organ traits derived from abdominal MRI using deep learning,” Elife, vol. 10, p. e65554, 2021.

[6] A. Y. Chu et al., “Multiethnic genome-wide meta-analysis of ectopic fat depots identifies loci associated with adipocyte development and differentiation,” Nature genetics, vol. 49, no. 1, pp. 125–130, 2017.

[7] G. R. Hunter, B. A. Gower, and B. L. Kane, “Age related shift in visceral fat,” International journal of body composition research, vol. 8, no. 3, p. 103, 2010.

[8] J. Li, Y. Xie, F. Yuan, B. Song, and C. Tang, “Noninvasive quantification of pancreatic fat in healthy male population using chemical shift magnetic resonance imaging: Effect of aging on pancreatic fat content,” Pancreas, vol. 40, no. 2, pp. 295–299, 2011.

[9] J. L. Kuk, T. J. Saunders, L. E. Davidson, and R. Ross, “Age-related changes in total and regional fat distribution,” Ageing research reviews, vol. 8, no. 4, pp. 339–348, 2009.

[10] S. Cheng et al., “Effect of aerobic exercise and diet on liver fat in pre-diabetic patients with non-alcoholic-fatty-liver-disease: A randomized controlled trial,” Scientific reports, vol. 7, no. 1, p. 15952, 2017.

[11] K. Young et al., “Influence of SNP* SNP interaction on BMI in e uropean a merican adolescents: Findings from the n ational l ongitudinal s tudy of a dolescent h ealth,” Pediatric obesity, vol. 11, no. 2, pp. 95–101, 2016.

[12] S. D’Silva, S. Chakraborty, and B. Kahali, “Concurrent outcomes from multiple approaches of epistasis analysis for human body mass index associated loci provide insights into obesity biology,” Scientific Reports, vol. 12, no. 1, p. 7306, 2022.

[13] E. Trépo, S. Romeo, J. Zucman-Rossi, and P. Nahon, “PNPLA3 gene in liver diseases,” Journal of hepatology, vol. 65, no. 2, pp. 399–412, 2016.

[14] J. Shen et al., “PNPLA 3 gene polymorphism and response to lifestyle modification in patients with nonalcoholic fatty liver disease,” Journal of gastroenterology and hepatology, vol. 30, no. 1, pp. 139–146, 2015.

[15] E. Herrera-Luis, K. Benke, H. Volk, C. Ladd-Acosta, and G. L. Wojcik, “Gene–environment interactions in human health,” Nature Reviews Genetics, vol. 25, no. 11, pp. 768–784, 2024.

[16] T. F. Mackay and R. R. Anholt, “Pleiotropy, epistasis and the genetic architecture of quantitative traits,” Nature Reviews Genetics, vol. 25, no. 9, pp. 639–657, 2024.

[17] X. Zhang and J. T. Bell, “Detecting genetic effects on phenotype variability to capture gene-by-environment interactions: A systematic method comparison,” G3: Genes, Genomes, Genetics, vol. 14, no. 4, p. jkae022, 2024.

[18] H. Wang et al., “Genotype-by-environment interactions inferred from genetic effects on phenotypic variability in the UK biobank,” Science advances, vol. 5, no. 8, p. eaaw3538, 2019.

[19] A. R. Marderstein, E. R. Davenport, S. Kulm, C. V. Van Hout, O. Elemento, and A. G. Clark, “Leveraging phenotypic variability to identify genetic interactions in human phenotypes,” The American Journal of Human Genetics, vol. 108, no. 1, pp. 49–67, 2021.

[20] J. Yang et al., “FTO genotype is associated with phenotypic variability of body mass index,” Nature, vol. 490, no. 7419, pp. 267–272, 2012.

[21] K. E. Westerman et al., “Variance-quantitative trait loci enable systematic discovery of gene-environment interactions for cardiometabolic serum biomarkers,” Nature Communications, vol. 13, no. 1, p. 3993, 2022.

[22] G. Shi, “Genome-wide variance quantitative trait locus analysis suggests small interaction effects in blood pressure traits,” Scientific Reports, vol. 12, no. 1, p. 12649, 2022.

[23] T. Lu, V. Forgetta, J. B. Richards, and C. M. Greenwood, “Genetic determinants of polygenic prediction accuracy within a population,” Genetics, vol. 222, no. 4, p. iyac158, 2022.

[24] J. Miao et al., “A quantile integral linear model to quantify genetic effects on phenotypic variability,” Proceedings of the National Academy of Sciences, vol. 119, no. 39, p. e2212959119, 2022.

[25] S. E. Petersen et al., “Imaging in population science: Cardiovascular magnetic resonance in 100,000 participants of UK biobank - rationale, challenges and approaches,” Journal of Cardiovascular Magnetic Resonance, vol. 15, no. 1, p. 46, 2013, doi: 10.1186/1532-429X-15-46.

[26] V. Groves et al., “How does variation in the body composition of both stimuli and participant modulate self-estimates of men’s body size?” Frontiers in Psychiatry, vol. 10, p. 720, 2019.

[27] J. Mbatchou et al., “Computationally efficient whole-genome regression for quantitative and binary traits,” Nature genetics, vol. 53, no. 7, pp. 1097–1103, 2021.

[28] F. Luo, F. Oldoni, and A. Das, “TM6SF2: A novel genetic player in nonalcoholic fatty liver and cardiovascular disease,” Hepatology communications, vol. 6, no. 3, pp. 448–460, 2022.

[29] M. M. Bos, R. Noordam, G. J. Blauw, P. E. Slagboom, P. C. Rensen, and D. van Heemst, “The ApoE 𝜀4 isoform: Can the risk of diseases be reduced by environmental factors?” The Journals of Gerontology: Series A, vol. 74, no. 1, pp. 99–107, 2019.

[30] N. Lan et al., “FTO–a common genetic basis for obesity and cancer,” Frontiers in genetics, vol. 11, p. 559138, 2020.

[31] X. Zhao et al., “FTO-dependent demethylation of N6-methyladenosine regulates mRNA splicing and is required for adipogenesis,” Cell research, vol. 24, no. 12, pp. 1403–1419, 2014.

[32] P. T. Williams, “Quantile-specific penetrance of genes affecting lipoproteins, adiposity and height,” PloS one, vol. 7, no. 1, p. e28764, 2012.

[33] B. L. Wajchenberg, “Subcutaneous and visceral adipose tissue: Their relation to the metabolic syndrome,” Endocrine reviews, vol. 21, no. 6, pp. 697–738, 2000.

[34] A. Nagral et al., “Gender differences in nonalcoholic fatty liver disease,” Euroasian journal of hepato-gastroenterology, vol. 12, no. Suppl 1, p. S19, 2022.

[35] G. Sveinbjornsson et al., “Multiomics study of nonalcoholic fatty liver disease,” Nature genetics, vol. 54, no. 11, pp. 1652–1663, 2022.

[36] J. Fu et al., “Effects of sex on the relationship between apolipoprotein e gene and serum lipid profiles in alzheimer’s disease,” Frontiers in Aging Neuroscience, vol. 14, p. 844066, 2022.

[37] F. K. Welty, A. H. Lichtenstein, P. H. R. Barrett, J. L. Jenner, G. G. Dolnikowski, and E. J. Schaefer, “Effects of ApoE genotype on ApoB-48 and ApoB-100 kinetics with stable isotopes in humans,” Arteriosclerosis, thrombosis, and vascular biology, vol. 20, no. 7, pp. 1807–1810, 2000.

[38] D. J. Sherman et al., “The fatty liver disease–causing protein PNPLA3-I148M alters lipid droplet–golgi dynamics,” Proceedings of the National Academy of Sciences, vol. 121, no. 18, p. e2318619121, 2024.

[39] W. G. Hill, M. E. Goddard, and P. M. Visscher, “Data and theory point to mainly additive genetic variance for complex traits,” PLoS genetics, vol. 4, no. 2, p. e1000008, 2008.

[40] V. Hivert et al., “Estimation of non-additive genetic variance in human complex traits from a large sample of unrelated individuals,” The American Journal of Human Genetics, vol. 108, no. 5, pp. 786–798, 2021.

[41] V. Nobili et al., “Influence of dietary pattern, physical activity, and I148M PNPLA3 on steatosis severity in at-risk adolescents,” Genes & nutrition, vol. 9, pp. 1–7, 2014.

[42] A. I. Young, F. Wauthier, and P. Donnelly, “Multiple novel gene-by-environment interactions modify the effect of FTO variants on body mass index,” Nature communications, vol. 7, no. 1, p. 12724, 2016.

[43] S. Gensluckner et al., “PNPLA3 and TM6SF2 exacerbate the impact of alcohol and metabolic dysfunction on liver fibrosis☆,” Jhep Reports, p. 101649, 2025.

[44] V. L. Chen et al., “Genetic risk accentuates dietary effects on hepatic steatosis, inflammation and fibrosis in a population-based cohort,” Journal of Hepatology, 2024.

[45] J. Boeckmans, A. Gatzios, J. M. Schattenberg, G. H. Koek, R. M. Rodrigues, and T. Vanhaecke, “PNPLA3 I148M and response to treatment for hepatic steatosis: A systematic review,” Liver International, vol. 43, no. 5, pp. 975–988, 2023.

[46] T. G. Richardson et al., “Characterising metabolomic signatures of lipid-modifying therapies through drug target mendelian randomisation,” PLoS biology, vol. 20, no. 2, p. e3001547, 2022.

[47] B.-T. Li et al., “Disruption of the ERLIN–TM6SF2–APOB complex destabilizes APOB and contributes to non-alcoholic fatty liver disease,” PLoS genetics, vol. 16, no. 8, p. e1008955, 2020.

[48] S. Prill et al., “The TM6SF2 E167K genetic variant induces lipid biosynthesis and reduces apolipoprotein b secretion in human hepatic 3D spheroids,” Scientific reports, vol. 9, no. 1, p. 11585, 2019.

[49] C. Bycroft et al., “The UK biobank resource with deep phenotyping and genomic data,” Nature, vol. 562, no. 7726, pp. 203–209, 2018.

[50] O. Pain, A. Al-Chalabi, and C. M. Lewis, “The GenoPred pipeline: A comprehensive and scalable pipeline for polygenic scoring,” Bioinformatics, vol. 40, no. 10, p. btae551, 2024.

[51] K. B. Hanscombe, J. R. Coleman, M. Traylor, and C. M. Lewis, “Ukbtools: An r package to manage and query UK biobank data,” PLoS One, vol. 14, no. 5, p. e0214311, 2019.

[52] F. Zhang et al., “OSCA: A tool for omic-data-based complex trait analysis,” Genome biology, vol. 20, pp. 1–13, 2019.

[53] H. Wang et al., “Genotype-by-environment interactions inferred from genetic effects on phenotypic variability in the UK biobank,” Science Advances, vol. 5, no. 8, p. eaaw3538, 2019, doi: 10.1126/sciadv.aaw3538.

[54] J. Yang et al., “Conditional and joint multiple-SNP analysis of GWAS summary statistics identifies additional variants influencing complex traits,” Nature genetics, vol. 44, no. 4, pp. 369–375, 2012.

[55] S. Purcell et al., “PLINK: A tool set for whole-genome association and population-based linkage analyses,” The American journal of human genetics, vol. 81, no. 3, pp. 559–575, 2007.

[56] L. A. Millard, N. M. Davies, T. R. Gaunt, G. Davey Smith, and K. Tilling, “Software application profile: PHESANT: A tool for performing automated phenome scans in UK biobank.” Oxford University Press, 2018.

[57] M. Higgins et al., “NHLBI family heart study: Objectives and design,” American journal of epidemiology, vol. 143, no. 12, pp. 1219–1228, 1996.

[58] M. F. Sinner et al., “Relation of circulating liver transaminase concentrations to risk of new-onset atrial fibrillation,” The American journal of cardiology, vol. 111, no. 2, pp. 219–224, 2013.

[59] M. B. A. L. 18. W. S. J. 18. W. V. A. 18. M. J. G. 18. C. M. S. 10. 18. Biobank and A. of Us Research Demonstration Project Teams Choi Seung Hoan 14 http://orcid.org/0000-0002-0322-8970 Wang Xin 14 http://orcid.org/0000-0001-6042-4487 Rosenthal Elisabeth A. 15, “Genomic data in the all of us research program,” Nature, vol. 627, no. 8003, pp. 340–346, 2024.

[60] S. Verdi et al., “TwinsUK: The UK adult twin registry update,” Twin Research and Human Genetics, vol. 22, no. 6, pp. 523–529, 2019.

[61] C. J. Willer, Y. Li, and G. R. Abecasis, “METAL: Fast and efficient meta-analysis of genomewide association scans,” Bioinformatics, vol. 26, no. 17, pp. 2190–2191, 2010.

