## Supplementary Materials for "Genetic Impacts on Variability of Body Fat Distribution Uncover Gene-Environment and Gene-Gene Interactions"

Supplementary Figure 1. Visualization of vQTLs and quantile QTLs. A) The phenotype distribution across the three genotype groups of each vQTL. B) The quantile QTL results of vQTLs. It represents the distributions of phenotypes in the first, fifth, and tenth quantile groups. The slope is the effect size of the genetic variant of each quantile group. In both plots, genotype 0 represents homozygous for the reference allele while genotype 2 represents homozygous for the alternative allele. The reference/alternative alleles are as follows: rs738408 (C/T), rs429358 (T/C), rs58542926 (C/T), and rs1285330517 (CAAA/C).

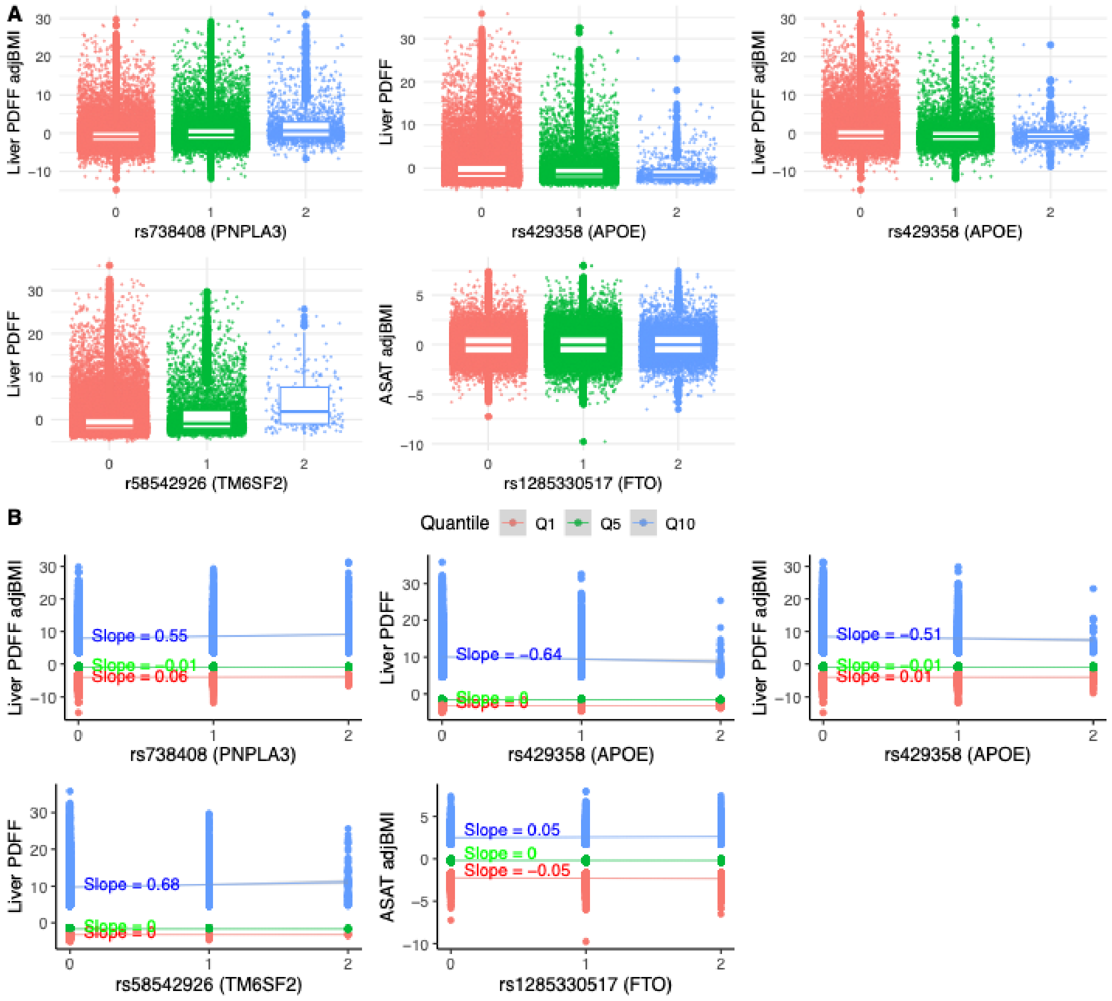

Supplementary Figure 2. vQTL-sex interaction effects.

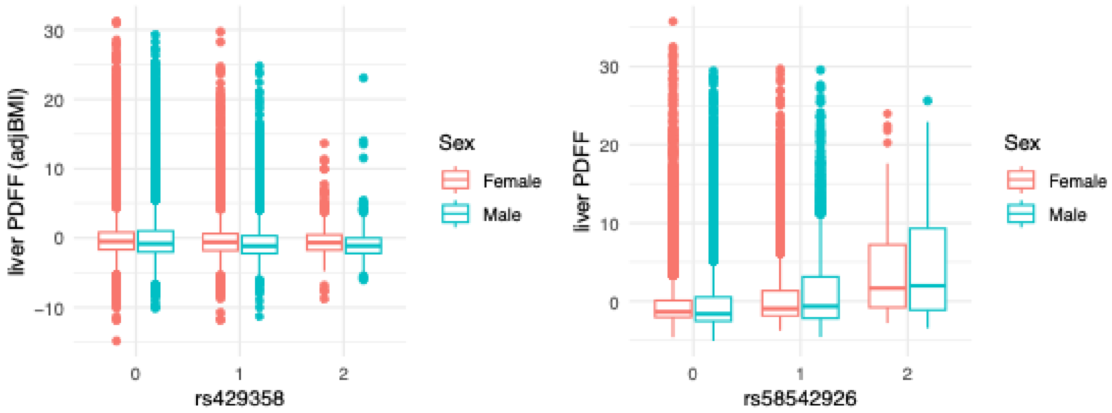

Supplementary Figure 3. Epistasis effects between rs58542926 and rs429358 on liver PDFF.

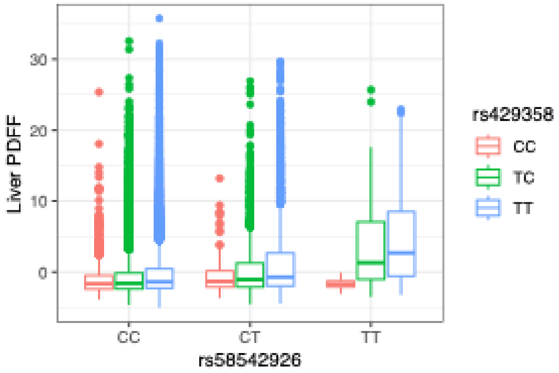

Supplementary Table 1. Summary statistics of quantile QTLs analysis.

| SNP | Phenotype | Quantile | Effect size | P-value |
|---|---|---|---|---|
| rs429358 | Liver PDFF | Q1 | 0.005 | 0.59 |
| rs429358 | Liver PDFF | Q5 | -0.003 | 0.37 |
| rs429358 | Liver PDFF | Q10 | -0.64 | 1e-04 |
| rs58542926 | Liver PDFF | Q1 | 0.002 | 0.88 |
| rs58542926 | Liver PDFF | Q5 | -0.0005 | 0.91 |
| rs58542926 | Liver PDFF | Q10 | 0.68 | 1e-05 |
| rs429358 | Liver PDFF adjBMI | Q1 | 0.01 | 0.76 |
| rs429358 | Liver PDFF adjBMI | Q5 | -0.006 | 0.11 |
| rs429358 | Liver PDFF adjBMI | Q10 | -0.51 | 2e-03 |
| rs738408 | Liver PDFF adjBMI | Q1 | 0.06 | 0.08 |
| rs738408 | Liver PDFF adjBMI | Q5 | -0.008 | 0.03 |
| rs738408 | Liver PDFF adjBMI | Q10 | 0.55 | 4.5e-07 |
| rs1285330517 | ASAT adjBMI | Q1 | -0.05 | 1e-04 |
| rs1285330517 | ASAT adjBMI | Q5 | -0.001 | 0.6 |
| rs1285330517 | ASAT adjBMI | Q10 | 0.05 | 3e-03 |

Supplementary Table 2. vQTL validation in datasets processed by other institutions

| vQTL | Phenotype | Field ID | P(BF) | P(DRM) | P(SVLM) |
|---|---|---|---|---|---|
| rs58542926 | Liver PDFF | 24352 | 2.77e-104 | 3.02e-102 | 5.64e-47 |
| rs58542926 | Liver PDFF | 21088 | 1.53e-70 | 3.29e-70 | 7.08e-35 |
| rs58542926 | Liver PDFF | 40061 | 1.05e-91 | 2.18e-89 | 2.15e-39 |
| rs429358 | Liver PDFF | 24352 | 1.28e-31 | 9.26e-33 | 7.74e-19 |
| rs429358 | Liver PDFF | 21088 | 4.56e-20 | 3.96e-21 | 1.37e-12 |
| rs429358 | Liver PDFF | 40061 | 2.65e-28 | 1.97e-29 | 3.83e-15 |
| rs429358 | Liver PDFF.adjBMI | 24352 | 4.62e-24 | 4.81e-25 | 8.08e-17 |
| rs429358 | Liver PDFF.adjBMI | 21088 | 3.46e-14 | 3.71e-15 | 3.58e-11 |
| rs429358 | Liver PDFF.adjBMI | 40061 | 4.38e-21 | 3.71e-22 | 5.55e-14 |
| rs738408 | Liver PDFF.adjBMI | 24352 | 2.31e-76 | 4.80e-75 | 2.93e-46 |
| rs738408 | Liver PDFF.adjBMI | 21088 | 1.35e-50 | 2.72e-49 | 7.84e-28 |
| rs738408 | Liver PDFF.adjBMI | 40061 | 3.22e-76 | 4.22e-73 | 9.04e-44 |
| rs1285330517 | ASAT.adjBMI | 22408 | 9.67e-09 | 2.23e-08 | 3.22e-10 |
| rs1285330517 | ASAT.adjBMI | 21086 | 1.43e-05 | 1.07e-05 | 2.44e-07 |

Supplementary Table 3. vQTL validations in individuals with follow-up visits. The P-values shown here is the largest P-values of vQTL identification methods. NS means not significant. For example, in the first row, "P<3e-02 in BF and DRM, NS in SVLM" means this vQTL was validated by BF and DRM methods and the larger P-value between these two methods was 3e-02, however, this vQTL was not validated by SVLM method.

| **Lead vQTL** | **MRI traits** | **Sample size** | **Discovery set** | **Follow-up set** |
|---|---|---|---|---|
| rs738408 | LiverPDFF.adjBMI | 2076 | P<4e-03 in BF, DRM, SVLM | P<3e-02 in BF and DRM, NS in SVLM |
| rs429358 | LiverPDFF.adjBMI | 2076 | NS in BF, DRM, and SVLM | NS in BF, DRM, and SVLM |
| rs429358 | LiverPDFF | 2210 | P<4e-02, NS in SVLM | NS in BF, DRM, and SVLM |
| rs58542926 | LiverPDFF | 2210 | P<9e-03 in BF, DRM, SVLM | P<7.8e-04 in BF, DRM, SVLM |
| rs1285330517 | ASAT.adjBMI | 2380 | NS in BF, DRM, SVLM | NS in BF, DRM, SVLM |

Supplementary Table 4: Power analysis using a subset of discovery samples.

| Sample size | Tested associations | BF | DRM | SVLM | Epistasis |
|---|---|---|---|---|---|
| 2000 | rs429358 - liver PDFF | 0.66 | 0.74 | 0.52 | NA |
| 2000 | rs429358 - liver PDFF adjBMI | 0.54 | 0.74 | 0.51 | NA |
| 2000 | rs58542926 - liver PDFF | 0.99 | 0.99 | 0.86 | NA |
| 2000 | rs738408 - liver PDFF adjBMI | 0.93 | 0.96 | 0.77 | NA |
| 2000 | rs1285330517 - ASAT adjBMI | 0.13 | 0.12 | 0.15 | NA |
| 2500 | rs429358 - liver PDFF | 0.73 | 0.8 | 0.59 | NA |
| 2500 | rs429358 - liver PDFF adjBMI | 0.72 | 0.84 | 0.71 | NA |
| 2500 | rs58542926 - liver PDFF | 1 | 0.99 | 0.94 | NA |
| 2500 | rs738408 - liver PDFF adjBMI | 0.99 | 1 | 0.95 | NA |
| 2500 | rs1285330517 - ASAT adjBMI | 0.26 | 0.24 | 0.31 | NA |
| 3000 | rs429358 - liver PDFF | 0.86 | 0.96 | 0.78 | NA |
| 3000 | rs429358 - liver PDFF adjBMI | 0.73 | 0.87 | 0.7 | NA |
| 3000 | rs58542926 - liver PDFF | 1 | 1 | 0.96 | NA |
| 3000 | rs738408 - liver PDFF adjBMI | 1 | 1 | 0.97 | NA |
| 3000 | rs1285330517 - ASAT adjBMI | 0.3 | 0.36 | 0.41 | NA |
| 2000 | Epistasis | NA | NA | NA | 0.14 |
| 2500 | Epistasis | NA | NA | NA | 0.18 |
| 3000 | Epistasis | NA | NA | NA | 0.2 |

Supplementary Table 5. Environmental factors and manually assigned categories. The Field ID and Category ID are both from the UK Biobank data resource.

| **Field ID** | **Description** | **Category ID** | **Category** |
|---|---|---|---|
| 1558 | Alcohol intake frequency | 100051 | alcohol consumption |
| 1568 | Average weekly red wine intake | 100051 | alcohol consumption |
| 1578 | Average weekly champagne plus white wine intake | 100051 | alcohol consumption |
| 1588 | Average weekly beer plus cider intake | 100051 | alcohol consumption |
| 1598 | Average weekly spirits intake | 100051 | alcohol consumption |
| 1608 | Average weekly fortified wine intake | 100051 | alcohol consumption |
| 5364 | Average weekly intake of other alcoholic drinks | 100051 | alcohol consumption |
| 21003 | Age when attended assessment centre | 1001 | demographic information |
| 31 | Sex | 1001 | demographic information |
| 21101 | Water hardness (WHO classification) | 603 | diet |
| 6155 | Vitamin and mineral supplements | 100045 | diet |
| 6179 | Mineral and other dietary supplements | 100045 | diet |
| 1548 | Variation in diet | 100052 | diet |
| 1329 | Oily fish intake | 100052 | diet |
| 1339 | Non-oily fish intake | 100052 | diet |
| 1349 | Processed meat intake | 100052 | diet |
| 1359 | Poultry intake | 100052 | diet |
| 1369 | Beef intake | 100052 | diet |
| 1379 | Lamb/mutton intake | 100052 | diet |
| 1389 | Pork intake | 100052 | diet |
| 1408 | Cheese intake | 100052 | diet |
| 1478 | Salt added to food | 100052 | diet |
| 1518 | Hot drink temperature | 100052 | diet |
| 1289 | Cooked vegetable intake | 100052 | diet |
| 1299 | Salad / raw vegetable intake | 100052 | diet |
| 1309 | Fresh fruit intake | 100052 | diet |
| 1319 | Dried fruit intake | 100052 | diet |
| 1438 | Bread intake | 100052 | diet |
| 1458 | Cereal intake | 100052 | diet |
| 1488 | Tea intake | 100052 | diet |
| 1498 | Coffee intake | 100052 | diet |
| 1528 | Water intake | 100052 | diet |
| 22032 | IPAQ activity group | 54 | physical activity |
| 22039 | MET minutes per week for vigorous activity | 54 | physical activity |
| 22035 | At or above moderate/vigorous recommendation | 54 | physical activity |
| 22036 | At or above moderate/vigorous/walking recommendation | 54 | physical activity |
| 22033 | Summed days activity | 54 | physical activity |
| 22034 | Summed minutes activity | 54 | physical activity |

| | | | |
|---|---|---|---|
| 22037 | MET minutes per week for walking | 54 | physical activity |
| 22038 | MET minutes per week for moderate activity | 54 | physical activity |
| 22040 | Summed MET minutes per week for all activity | 54 | physical activity |
| 1110 | Length of mobile phone use | 100053 | sedentary behavior |
| 924 | Usual walking pace | 100054 | sedentary behavior |
| 1070 | Time spent watching television (TV) | 100054 | sedentary behavior |
| 1080 | Time spent using computer | 100054 | sedentary behavior |
| 1090 | Time spent driving | 100054 | sedentary behavior |
| 864 | Number of days/week walked 10+ minutes | 100054 | sedentary behavior |
| 874 | Duration of walks | 100054 | sedentary behavior |
| 884 | Number of days/week of moderate physical activity 10+ minutes | 100054 | sedentary behavior |
| 894 | Duration of moderate activity | 100054 | sedentary behavior |
| 904 | Number of days/week of vigorous physical activity 10+ minutes | 100054 | sedentary behavior |
| 1180 | Morning/evening person (chronotype) | 100057 | sleep |
| 1190 | Nap during day | 100057 | sleep |
| 1200 | Sleeplessness / insomnia | 100057 | sleep |
| 1160 | Sleep duration | 100057 | sleep |
| 20116 | Smoking status | 100058 | smoking |
| 1239 | Current tobacco smoking | 100058 | smoking |

Supplementary Table 6. vQTL-environment interactions.

| vQTL | MRI trait | Environmental factor | Sample size | P(GE) |
|---|---|---|---|---|
| **Individual variables** | | | | |
| rs738408 | Liver PDFF.adjBMI | Variation in diet | 39,843 | 4.37E-06 |
| rs738408 | Liver PDFF adjBMI | Usual walking pace | 39,835 | 2.69E-06 |
| rs738408 | Liver PDFF.adjBMI | Time spent watching television (TV) | 39,772 | 1.46E-09 |
| rs738408 | Liver PDFF.adjBMI | Time spent driving | 39,529 | 1.13E-04 |
| rs429358 | Liver PDFF | MET minutes per week for vigorous activity | 35,231 | 1.10E-04 |
| rs429358 | Liver PDFF | Usual walking pace | 41,195 | 1.71E-04 |
| rs58542926 | Liver PDFF | Average weekly red wine intake | 29,872 | 6.19E-06 |
| rs58542926 | Liver PDFF | Time spent using computer | 41,120 | 5.68E-06 |
| rs1285330517 | ASAT.adjBMI | Nap during day | 47,452 | 1.94E-04 |
| rs429358 | Liver PDFF.adjBMI | Sex | 40,144 | 5.61E-05 |
| rs58542926 | Liver PDFF | Sex | 41,534 | 2.78E-07 |
| **Category PCs** | | | | |
| rs58542926 | Liver PDFF | alcohol_PC2 | 29,694 | 4.86E-08 |
| rs738408 | Liver PDFF.adjBMI | alcohol_PC2 | 28,742 | 1.68E-05 |
| **Whole group PC** | | | | |
| rs58542926 | Liver PDFF | PC2 | 17,297 | 5.30E-06 |
| rs738408 | Liver PDFF.adjBMI | PC2 | 16,735 | 2.37E-05 |
| rs429358 | Liver PDFF.adjBMI | PC1 | 16,746 | 0.001262313 |
| rs429358 | Liver PDFF.adjBMI | PC2 | 16,746 | 0.001697712 |

Supplementary Table 7. vQTL-sex-environment interactions.

| vQTL | Environmental factor | trait | N | P_G-sex-E |
|---|---|---|---|---|
| rs58542926 | Length of mobile phone use | Liver PDFF | 40915 | 1.26E-05 |
| rs58542926 | Morning/evening person (chronotype) | Liver PDFF | 37533 | 1.59E-05 |
| rs429358 | Sleeplessness / insomnia | Liver PDFF adjBMI | 39886 | 2.64E-05 |
| rs429358 | Sleeplessness / insomnia | Liver PDFF | 41222 | 0.00010101 |
| rs58542926 | Oily fish intake | Liver PDFF | 41201 | 1.90E-05 |
| rs58542926 | Non-oily fish intake | Liver PDFF | 41182 | 0.00011486 |
| rs1285330517 | Processed meat intake | ASAT adjBMI | 47425 | 2.84E-08 |
| rs58542926 | Poultry intake | Liver PDFF | 41229 | 3.13E-05 |
| rs1285330517 | Poultry intake | ASAT adjBMI | 47426 | 6.20E-06 |
| rs1285330517 | Beef intake | ASAT adjBMI | 47380 | 2.00E-05 |
| rs58542926 | Cheese intake | Liver PDFF | 40346 | 2.96E-06 |
| rs429358 | Cheese intake | Liver PDFF adjBMI | 39039 | 4.03E-05 |
| rs429358 | Hot drink temperature | Liver PDFF adjBMI | 39533 | 2.94E-06 |
| rs58542926 | Hot drink temperature | Liver PDFF | 40857 | 5.33E-08 |
| rs58542926 | Alcohol intake frequency. | Liver PDFF | 41238 | 5.82E-07 |
| rs58542926 | Average weekly champagne plus white wine intake | Liver PDFF | 29857 | 6.22E-05 |
| rs1285330517 | Average weekly beer plus cider intake | ASAT adjBMI | 34415 | 0.00014038 |
| rs1285330517 | Average weekly spirits intake | ASAT adjBMI | 34363 | 1.82E-06 |
| rs1285330517 | Smoking status | ASAT adjBMI | 47280 | 4.39E-05 |
| rs58542926 | Age when attended assessment centre | Liver PDFF | 41534 | 1.71E-06 |
| rs429358 | Age when attended assessment centre | Liver PDFF adjBMI | 40144 | 9.28E-06 |
| rs429358 | Age when attended assessment centre | Liver PDFF | 41534 | 0.0001715 |
| rs429358 | IPAQ activity group | Liver PDFF adjBMI | 34097 | 1.03E-05 |
| rs58542926 | IPAQ activity group | Liver PDFF | 35231 | 3.71E-06 |
| rs429358 | At or above moderate/vigorous recommendation | Liver PDFF adjBMI | 34097 | 0.00017641 |

| | | | | |
|---|---|---|---|---|
| rs58542926 | At or above moderate/vigorous/walking recommendation | Liver PDFF | 35231 | 1.12E-07 |
| rs58542926 | MET minutes per week for vigorous activity | Liver PDFF | 35231 | 3.06E-07 |
| rs429358 | MET minutes per week for vigorous activity | Liver PDFF | 35231 | 1.16E-05 |
| rs429358 | MET minutes per week for vigorous activity | Liver PDFF adjBMI | 34097 | 1.85E-07 |
| rs429358 | Usual walking pace | Liver PDFF adjBMI | 39864 | 1.30E-05 |
| rs58542926 | Usual walking pace | Liver PDFF | 41195 | 1.09E-11 |

Supplementary Table 8. vQTL validation in liver serum markers.

| vQTL | Liver Marker | Sample Size | P(BF) | P(DRM) | P(SVLM) |
|---|---|---|---|---|---|
| rs429358 | ALT | 362863 | 6.2e-35 | 1.2e-35 | 1.1e-05 |
| rs58542926 | ALT | 362863 | 4.2e-72 | 3.2e-72 | 3.9e-10 |
| rs738408 | ALT | 362863 | 8.4e-314 | 6.6e-297 | 4e-31 |
| rs429358 | ALT.adjBMI | 361697 | 8.9e-27 | 2.2e-27 | 5.8e-05 |
| rs58542926 | ALT.adjBMI | 361697 | 1.1e-61 | 5.7e-62 | 5.4e-09 |
| rs738408 | ALT.adjBMI | 361697 | 2.3e-248 | 2e-233 | 1.2e-24 |
| rs429358 | AST | 361651 | 8.2e-06 | 2.1e-06 | 0.5 |
| rs58542926 | AST | 361651 | 6.1e-27 | 7.1e-25 | 0.2 |
| rs738408 | AST | 361651 | 1.5e-132 | 1.3e-126 | 9.6e-06 |
| rs429358 | AST.adjBMI | 360490 | 8.1e-05 | 2.5e-05 | 0.5 |
| rs58542926 | AST.adjBMI | 360490 | 1.5e-24 | 1.5e-22 | 0.2 |
| rs738408 | AST.adjBMI | 360490 | 3.9e-115 | 6.1e-110 | 2e-05 |
| rs429358 | ALP | 363014 | 0.002 | 0.0005 | 0.3 |
| rs58542926 | ALP | 363014 | 2.5e-06 | 4.3e-07 | 0.04 |
| rs738408 | ALP | 363014 | 0.01 | 0.005 | 0.6 |
| rs429358 | ALP.adjBMI | 361845 | 0.0009 | 0.0002 | 0.3 |
| rs58542926 | ALP.adjBMI | 361845 | 3.2e-05 | 5.9e-06 | 0.05 |
| rs738408 | ALP.adjBMI | 361845 | 0.06 | 0.02 | 0.6 |
| rs429358 | Albumin | 332464 | 0.05 | 0.02 | 0.2 |
| rs58542926 | Albumin | 332464 | 0.07 | 0.03 | 0.01 |
| rs738408 | Albumin | 332464 | 0.5 | 0.27 | 0.02 |
| rs429358 | Albumin.adjBMI | 331383 | 0.07 | 0.02 | 0.2 |
| rs58542926 | Albumin.adjBMI | 331383 | 0.06 | 0.02 | 0.006 |
| rs738408 | Albumin.adjBMI | 331383 | 0.3 | 0.1 | 0.01 |
| rs429358 | DirectBilirubin | 308944 | 0.004 | 0.001 | 0.09 |
| rs58542926 | DirectBilirubin | 308944 | 5.6e-08 | 4.2e-07 | 0.7 |
| rs738408 | DirectBilirubin | 308944 | 0.004 | 0.003 | 0.5 |
| rs429358 | DirectBilirubin.adjBMI | 308014 | 0.003 | 0.0008 | 0.09 |
| rs58542926 | DirectBilirubin.adjBMI | 308014 | 4.6e-08 | 2.9e-07 | 0.7 |
| rs738408 | DirectBilirubin.adjBMI | 308014 | 0.001 | 0.0008 | 0.5 |
| rs429358 | GGT | 362815 | 1.7e-05 | 6.4e-06 | 0.04 |
| rs58542926 | GGT | 362815 | 0.0002 | 0.0005 | 0.07 |
| rs738408 | GGT | 362815 | 0.5 | 0.2 | 0.05 |
| rs429358 | GGT.adjBMI | 361648 | 0.0001 | 4.9e-05 | 0.05 |
| rs58542926 | GGT.adjBMI | 361648 | 0.0005 | 0.002 | 0.08 |
| rs738408 | GGT.adjBMI | 361648 | 0.7 | 0.4 | 0.07 |
| rs429358 | APOB | 361221 | 1.6e-56 | 9e-58 | 1.8e-49 |
| rs58542926 | APOB | 361221 | 5.7e-08 | 7.7e-09 | 0.001 |
| rs738408 | APOB | 361221 | 0.7 | 0.4 | 0.4 |

| | | | | | |
|---|---|---|---|---|---|
| rs429358 | APOB.adjBMI | 360059 | 3e-54 | 2e-55 | 1.1e-47 |
| rs58542926 | APOB.adjBMI | 360059 | 4.4e-06 | 6.8e-07 | 0.006 |
| rs738408 | APOB.adjBMI | 360059 | 0.8 | 0.6 | 0.6 |

Supplementary Table 9. Validation of vQTL-environment interaction effects on liver markers.

| vQTL | Environmental Factor | Liver Marker | N | $P_{GE}$ |
|---|---|---|---|---|
| rs58542926 | Sex | ALT.adjBMI | 361870 | 1.53e-07 |
| rs58542926 | Sex | ALT | 363037 | 8.90e-07 |
| rs58542926 | Sex | AST | 361825 | 9.20e-05 |
| rs58542926 | Sex | AST.adjBMI | 360663 | 5.29e-05 |
| rs58542926 | Sex | APOB.adjBMI | 360231 | 5.94e-25 |
| rs58542926 | Sex | APOB | 361394 | 6.37e-25 |
| rs429358 | MET minutes per week for vigorous activity | GGT.adjBMI | 282680 | 5.63e-03 |
| rs429358 | Sex | APOB | 361394 | 2.82e-16 |
| rs429358 | Sex | APOB.adjBMI | 360231 | 7.13e-17 |
| rs738408 | Time spent watching television (TV) | ALT | 343173 | 4.67e-11 |
| rs738408 | Time spent watching television (TV) | ALT.adjBMI | 342086 | 5.24e-11 |
| rs738408 | Time spent watching television (TV) | AST | 342020 | 3.42e-19 |
| rs738408 | Time spent watching television (TV) | AST.adjBMI | 340938 | 8.00e-19 |
| rs738408 | Time spent driving | ALT | 242686 | 8.51e-09 |
| rs738408 | Time spent driving | AST.adjBMI | 240967 | 3.44e-04 |
| rs738408 | Time spent driving | ALT.adjBMI | 241797 | 1.03e-09 |
| rs738408 | Variation in diet | ALT | 361711 | 6.61e-08 |
| rs738408 | Variation in diet | AST.adjBMI | 359372 | 6.71e-08 |
| rs738408 | Variation in diet | AST | 360505 | 2.39e-08 |
| rs738408 | Variation in diet | ALT.adjBMI | 360573 | 7.12e-07 |
| rs738408 | Usual walking pace | ALT | 360629 | 1.24e-10 |
| rs738408 | Usual walking pace | AST.adjBMI | 358593 | 1.31e-20 |
| rs738408 | Usual walking pace | AST | 359425 | 1.40e-19 |
| rs738408 | Usual walking pace | ALT.adjBMI | 359795 | 6.29e-13 |

Supplementary Table 10. vQTL replication on CT-measured liver fat from Framingham Heart Study. The replication was conducted using BF, DRM and SVLM methods, and the P-value listed in the table shows the largest observed significant P-value. For example, P<7.5e-19 means the largest significant P-value observed from all three vQTL detection methods is 7.5e-19. NS means not significant.

| ***Liver fat (without BMI adjustment)*** | | |
|---|---|---|
| | Discovery set (N = 40,145-41,535) | Replication set (N = 2,794) |
| rs738408 | NS | P<1.2e-08 |
| rs429358 | P<7.5e-19 | NS |
| rs58542926 | P<4.3e-17 | P<4.8e-03 |
| rs58542926*rs429358 | 3.6e-04 | NS |
| ***Liver fat (with BMI adjustment)*** | | |
| | Discovery set (N = 40,145-41,535) | Replication set (N = 2,794) |
| rs738408 | P<2.3e-46 | P<2.3e-07 |
| rs429358 | P<7.8e-17 | NS |
| rs58542926 | NS | P<1.8e-03 |
| rs58542926*rs429358 | 2.2e-05 | NS |

Supplementary Table 11. Replication results on liver function markers with and without BMI adjustment. The replication was conducted using BF, DRM and SVLM methods, and the threshold shows the largest observed significant P-value. NS means not significant. For example, "P<0.011, NS in SVLM" means the largest significant P-value observed from all vQTL detection methods is 0.011, but this vQTL was not detected by SVLM method.

| | | All of Us | | Framingham Heart Study | | TwinsUK | | UKB_exclude | |
|---|---|---|---|---|---|---|---|---|---|
| vQTL | Liver markers | Sample size | P-value | Sample size | P-value | Sample size | P-value | Sample size | P-value |
| rs738408 | ALT | 44,358 | P<0.011, NS in SVLM | 5,570 | P<0.01 | 2,837 | P<0.02, NS in DRM and SVLM | 323,345 | P<2.6e-31 |
| rs738408 | AST | 49,961 | NS | 5,570 | P<2e-3, NS in SVLM | 1,243 | NS | 322,278 | P<9.3e-5 |
| rs429358 | ALT | 44,360 | NS | 5,570 | NS | 2,837 | P<0.03, NS in DRM and BF | 323,345 | P<2e-5 |
| rs429358 | AST | 49,962 | NS | 5,570 | NS | 1,243 | NS | 322,278 | P<6.1e-5, NS in SVLM |
| rs58542926 | ALT | 44,363 | NS | 5,570 | NS | 2,837 | NS | 323,345 | P<6e-7 |
| rs58542926 | AST | 49,966 | NS | 5,570 | NS | 1,243 | P<0.04, NS in BF | 322,278 | P<3.6e-21, NS in SVLM |
| rs58542926* rs429358 | ALT | 44,363 | NS | 5,570 | NS | 2,837 | P=0.03 | 323,345 | P=2.6e-5 |
| rs58542926* rs429358 | AST | 49,966 | NS | 5,570 | NS | 1,243 | NS | 322,278 | P=2.5e-3 |
| rs738408 | ALT.adjBMI | 44,358 | P< 0.011, NS in SVLM | 5,563 | P<0.03 | 2,376 | NS | 322,217 | P<3.7e-35 |
| rs738408 | AST.adjBMI | 49,961 | NS | 5,563 | P<3e-3, NS in SVLM | 786 | NS | 321,155 | P<1.8e-5 |
| rs429358 | ALT.adjBMI | 44,360 | NS | 5,563 | NS | 2,376 | P<0.03, NS in BF | 322,217 | P<8.8e-5 |
| rs429358 | AST.adjBMI | 49,962 | NS | 5,563 | NS | 786 | NS | 321,155 | P<4e-4, NS in SVLM |
| rs58542926 | ALT.adjBMI | 44,363 | NS | 5,563 | NS | 2,376 | NS | 322,217 | P<4.6e-6 |
| rs58542926 | AST.adjBMI | 49,966 | NS | 5,563 | NS | 786 | NS | 321,155 | P<4.5e-19, NS in SVLM |
| rs58542926* rs429358 | ALT.adjBMI | 44,363 | NS | 5,563 | NS | 2,376 | P=0.01 | 322,217 | P=1.4e-5 |
| rs58542926* rs429358 | AST.adjBMI | 49,966 | NS | 5,563 | NS | 786 | NS | 321,155 | P=2.6e-3 |

Supplementary Table 12. The significant (P<.05/3) meta-analysis results of vQTL effects on liver markers.

| vQTL | Liver Markers | Sample Size | P-value | Summary Statistics | Cohort |
|---|---|---|---|---|---|
| rs429358 | ALT.adjBMI | 374516 | 1.28e-23 | BF | All of Us, FHS, TwinsUK, UKB_exclude |
| rs429358 | ALT.adjBMI | 374516 | 5.82e-21 | DRM | All of Us, FHS, TwinsUK, UKB_exclude |
| rs429358 | ALT.adjBMI | 374516 | 9.78e-05 | SVLM | All of Us, FHS, TwinsUK, UKB_exclude |
| rs429358 | ALT | 376112 | 1.42e-29 | BF | All of Us, FHS, TwinsUK, UKB_exclude |
| rs429358 | ALT | 376112 | 7.69e-27 | DRM | All of Us, FHS, TwinsUK, UKB_exclude |
| rs429358 | ALT | 376112 | 2.40e-05 | SVLM | All of Us, FHS, TwinsUK, UKB_exclude |
| rs429358 | AST.adjBMI | 377466 | 2.82e-04 | BF | All of Us, FHS, TwinsUK, UKB_exclude |
| rs429358 | AST.adjBMI | 377466 | 1.78e-03 | DRM | All of Us, FHS, TwinsUK, UKB_exclude |
| rs429358 | AST | 379053 | 4.08e-05 | BF | All of Us, FHS, TwinsUK, UKB_exclude |
| rs429358 | AST | 379053 | 3.54e-04 | DRM | All of Us, FHS, TwinsUK, UKB_exclude |
| rs58542926 | ALT.adjBMI | 374519 | 4.56e-48 | BF | All of Us, FHS, TwinsUK, UKB_exclude |
| rs58542926 | ALT.adjBMI | 374519 | 9.51e-42 | DRM | All of Us, FHS, TwinsUK, UKB_exclude |
| rs58542926 | ALT.adjBMI | 374519 | 5.52e-06 | SVLM | All of Us, FHS, TwinsUK, UKB_exclude |
| rs58542926 | ALT | 376115 | 1.38e-56 | BF | All of Us, FHS, TwinsUK, UKB_exclude |
| rs58542926 | ALT | 376115 | 1.66e-49 | DRM | All of Us, FHS, TwinsUK, UKB_exclude |
| rs58542926 | ALT | 376115 | 7.36e-07 | SVLM | All of Us, FHS, TwinsUK, UKB_exclude |
| rs58542926 | AST.adjBMI | 377470 | 1.37e-18 | BF | All of Us, FHS, TwinsUK, UKB_exclude |
| rs58542926 | AST.adjBMI | 377470 | 3.99e-16 | DRM | All of Us, FHS, TwinsUK, UKB_exclude |
| rs58542926 | AST | 379057 | 1.15e-20 | BF | All of Us, FHS, TwinsUK, UKB_exclude |
| rs58542926 | AST | 379057 | 1.47e-17 | DRM | All of Us, FHS, TwinsUK, UKB_exclude |
| rs738408 | ALT | 376110 | 4.58e-277 | BF | All of Us, FHS, TwinsUK, UKB_exclude |
| rs738408 | ALT.adjBMI | 374514 | 6.26e-221 | BF | All of Us, FHS, TwinsUK, UKB_exclude |
| rs738408 | ALT.adjBMI | 374514 | 2.92e-171 | DRM | All of Us, FHS, TwinsUK, UKB_exclude |
| rs738408 | ALT.adjBMI | 52297 | 4.30e-05 | BF | All of Us, FHS, TwinsUK |
| rs738408 | ALT.adjBMI | 52297 | 1.68e-04 | DRM | All of Us, FHS, TwinsUK |
| rs738408 | ALT.adjBMI | 56310 | 1.46e-02 | DRM | All of Us, FHS, TwinsUK |
| rs738408 | ALT.adjBMI | 374514 | 5.78e-25 | SVLM | All of Us, FHS, TwinsUK, UKB_exclude |
| rs738408 | ALT | 376110 | 4.37e-218 | DRM | All of Us, FHS, TwinsUK, UKB_exclude |
| rs738408 | ALT | 376110 | 9.03e-31 | SVLM | All of Us, FHS, TwinsUK, UKB_exclude |
| rs738408 | ALT | 52765 | 1.15e-05 | BF | All of Us, FHS, TwinsUK |
| rs738408 | ALT | 52765 | 5.60e-05 | DRM | All of Us, FHS, TwinsUK |
| rs738408 | AST.adjBMI | 377465 | 4.83e-101 | BF | All of Us, FHS, TwinsUK, UKB_exclude |
| rs738408 | AST.adjBMI | 377465 | 8.04e-80 | DRM | All of Us, FHS, TwinsUK, UKB_exclude |
| rs738408 | AST.adjBMI | 377465 | 3.52e-06 | SVLM | All of Us, FHS, TwinsUK, UKB_exclude |
| rs738408 | AST.adjBMI | 56310 | 1.06e-02 | BF | All of Us, FHS, TwinsUK |
| rs738408 | AST | 379052 | 4.74e-115 | BF | All of Us, FHS, TwinsUK, UKB_exclude |
| rs738408 | AST | 379052 | 8.20e-92 | DRM | All of Us, FHS, TwinsUK, UKB_exclude |
| rs738408 | AST | 379052 | 1.94e-06 | SVLM | All of Us, FHS, TwinsUK, UKB_exclude |

| | | | | | |
|---|---|---|---|---|---|
| rs738408 | AST | 56774 | 1.14e-02 | BF | All of Us, FHS, TwinsUK |
| rs738408 | AST | 56774 | 1.20e-02 | DRM | All of Us, FHS, TwinsUK |
| rs58542926*rs429358 | AST.adjBMI | 377465 | 3.00e-03 | Epistasis | All of Us, FHS, TwinsUK, UKB_exclude |
| rs58542926*rs429358 | AST | 379052 | 3.20e-03 | Epistasis | All of Us, FHS, TwinsUK, UKB_exclude |
| rs58542926*rs429358 | ALT.adjBMI | 374516 | 3.20e-05 | Epistasis | All of Us, FHS, TwinsUK, UKB_exclude |
| rs58542926*rs429358 | AST | 376112 | 6.40e-05 | Epistasis | All of Us, FHS, TwinsUK, UKB_exclude |
